# Voyager: exploratory single-cell genomics data analysis with geospatial statistics

**DOI:** 10.1101/2023.07.20.549945

**Authors:** Lambda Moses, Pétur Helgi Einarsson, Kayla Jackson, Laura Luebbert, A. Sina Booeshaghi, Sindri Antonsson, Nicolas Bray, Páll Melsted, Lior Pachter

## Abstract

Exploratory spatial data analysis (ESDA) can be a powerful approach to understanding single-cell genomics datasets, but it is not yet part of standard data analysis workflows. In particular, geospatial analyses, which have been developed and refined for decades, have yet to be fully adapted and applied to spatial single-cell analysis. We introduce the Voyager platform, which systematically brings the geospatial ESDA tradition to (spatial) -omics, with local, bivariate, and multivariate spatial methods not yet commonly applied to spatial -omics, united by a uniform user interface. Using Voyager, we showcase biological insights that can be derived with its methods, such as biologically relevant negative spatial autocorrelation. Underlying Voyager is the SpatialFeatureExperiment data structure, which combines Simple Feature with SingleCellExperiment and AnnData to represent and operate on geometries bundled with gene expression data. Voyager has comprehensive tutorials demonstrating ESDA built on GitHub Actions to ensure reproducibility and scalability, using data from popular commercial technologies. Voyager is implemented in both R/Bioconductor and Python/PyPI, and features compatibility tests to ensure that both implementations return consistent results.

## Introduction

From the developing embryo to the hepatic lobule, spatial organization of cells is essential to the functions of many tissues. Recent breakthroughs in technology development, data collection, and data analysis tools for spatial transcriptomics, have led to a plethora of possibilities and applications^1^. Among these data analysis tools are overarching data analysis frameworks for data organization and exploratory data analysis (EDA), including frameworks such as Seurat^2^, squidpy^3^, Giotto^4^, and semla (formerly STUtility^5^), which add spatial data analysis functionalities and visualizations to traditional single-cell RNA-seq (scRNA-seq) analysis workflows. In addition, for the purpose of EDA, many visualization tools have been developed for spatial -omics data. Many of these visualization tools are designed to be scalable and interactive for large imaging-based data such as MERFISH^6^ and imaging mass cytometry, plotting gene expression and cell metadata in space, or utilizing virtual reality^7–11^.

EDA is an approach to understanding data “without many preconceived ideas, theories, or hypotheses”^12^. It encourages a mindset of asking questions and exploring possible answers by visualizing, transforming, and modeling relevant data, leading to further, more refined questions without a formal process or strict set of rules^13^. The spirit of EDA is that “it is important to understand what you *can do* before you learn to measure how *well* you seem to have *done* it”^14^. Exploratory *spatial* data analysis (ESDA) is EDA explicitly focusing on spatial aspects of the data, especially spatial autocorrelation, where nearby observations are not independent from each other^12^. ESDA has a long history of use in geography, where a rich tradition has been developed^15^. The widely used spatial autocorrelation metrics Moran’s I^16^ and Geary’s C^17^ are among the global univariate spatial statistics used in ESDA, which produce one set of results for the entire dataset. The characteristics of Moran’s I and Geary’s C have been further elaborated over the years^18–20^. In addition, there are tools to explore the length scale of spatial autocorrelation, such as the correlogram^21^ and variogram^22^. Local versions of spatial statistics, such as local Moran’s I^23^ and Getis-Ord Gi* ^24^ can be used to explore local spatial heterogeneity and find spatial clusters of high or low values, producing a set of results for each location. There are also spatially informed global and local bivariate and multivariate statistics that account for spatial autocorrelation and correlation between features simultaneously, such as Lee’s L^25^ and MULTISPATI PCA^26^.

Much of this ESDA tradition can benefit spatial -omics, but has not been utilized in existing EDA frameworks. For example, while Seurat, squidpy, Giotto, and semla implement some spatial analysis methods; none of these tools have systematically and comprehensively implemented the entire breadth of the geospatial ESDA tradition (Supplementary Table 1). In particular, methods to explore length scales and local spatial heterogeneity of spatial autocorrelation and spatially informed correlation among genes have not been fully explored in these existing frameworks (Supplementary Table 1). Furthermore, the data structures underlying Seurat, squidpy, Giotto, and semla have limited if any support for representation and operations on geometries such as cell segmentation polygons (Supplementary Table 2). In addition, packages that specialize in visualization often do not implement any ESDA methods. Many other spatial -omics data analysis packages focus on data preprocessing before the exploratory stage, such as image processing^27^, cell type deconvolution of Visium spots^28^, integrating data from different modalities or tissue sections^29^, and predicting gene expression from H&E images^30^.

Downstream analysis packages tend to implement novel methods for specific tasks such as finding spatially variable genes^31^, cell-cell interactions^32^, spatially informed dimension reduction^33^, and finding spatial regions defined by gene expression^34^, accompanied by claims of superior performance compared to existing packages performing the same tasks. These packages focus on “how *well* you seem to have *done* it” in Tukey’s words while “what you *can do*” remains unaddressed.

The Voyager project fills this gap by facilitating geospatial ESDA to spatial -omics, placing the spatial information front and center in the EDA workflow. The SpatialFeatureExperiment (SFE) object underlying Voyager brings geospatial Simple Features to spatial -omics, thereby enabling geometric operations, such as to relate characteristics of cells to those of Visium spots. EDA is to “feel free to investigate every idea that occurs to you"^13^. Many univariate, bivariate, and multivariate global and local methods are included (Supplementary Table 1), with a consistent user interface extensible to new methods, enabling researchers to easily investigate more such ideas. Some classic methods have been reimplemented to be more scalable to larger datasets, and we demonstrate the utility of ESDA by applying Voyager to real data. Our case studies show what we “can do” with an expanded palette of ESDA tools beyond areas commonly addressed by specialized downstream methods: we find correlation between library size and histological characteristics, and show an example of biologically relevant negative spatial autocorrelation. We also show that ESDA can be applied to non-spatial scRNA-seq datasets via the k-nearest-neighbor graph, thereby making Voyager immediately applicable to a wide range of single-cell genomics datasets.

Finally, we address an increasingly challenging problem in single-cell genomics, namely the divergence between R and Python^1^ implementations of standard methods, that has led to programming-language dependent results. Voyager is implemented in both R and Python, with compatibility tests to ensure that the two implementations give consistent results for core functionalities. Voyager also has comprehensive documentation and tutorials for common spatial -omics technologies and spatial analysis methods. While the tutorials are built on spatial transcriptomics, proteomics, and non-spatial scRNA-seq data, in principle Voyager can be applied to other -omics as well. The R packages Voyager, SpatialFeatureExperiment, and SFEData (which provides example datasets for the tutorials) are available on Bioconductor. The Python implementation is available on PyPI.

## Results

### The Voyager framework

Voyager is built on the SFE data structure, which bundles geometries such as cell segmentation polygons with gene expression data, and implements geometric operations on such geometries and related SFE data. Geometries in SFE are represented with Simple Features, which is a standard framework to represent geometries in the geospatial field and provides access to the GEOS C++ library for fast geometric operations. SFE extends existing scRNA-seq data structures and conforms to their syntaxes and conventions. In Python, AnnData^35^ is extended with GeoPandas^36^, and in R, SingleCellExperiment (SCE)^37^ and SpatialExperiment (SPE)^38^ are extended with sf^39^ for the Simple Features (Figure 1, Supplementary Note 1). Importantly, spatial information need not be physical: single-cell RNA-seq data structures can also be analyzed spatially, where “space” is abstracted to the k-nearest-neighbor graph.

**Figure 1:**
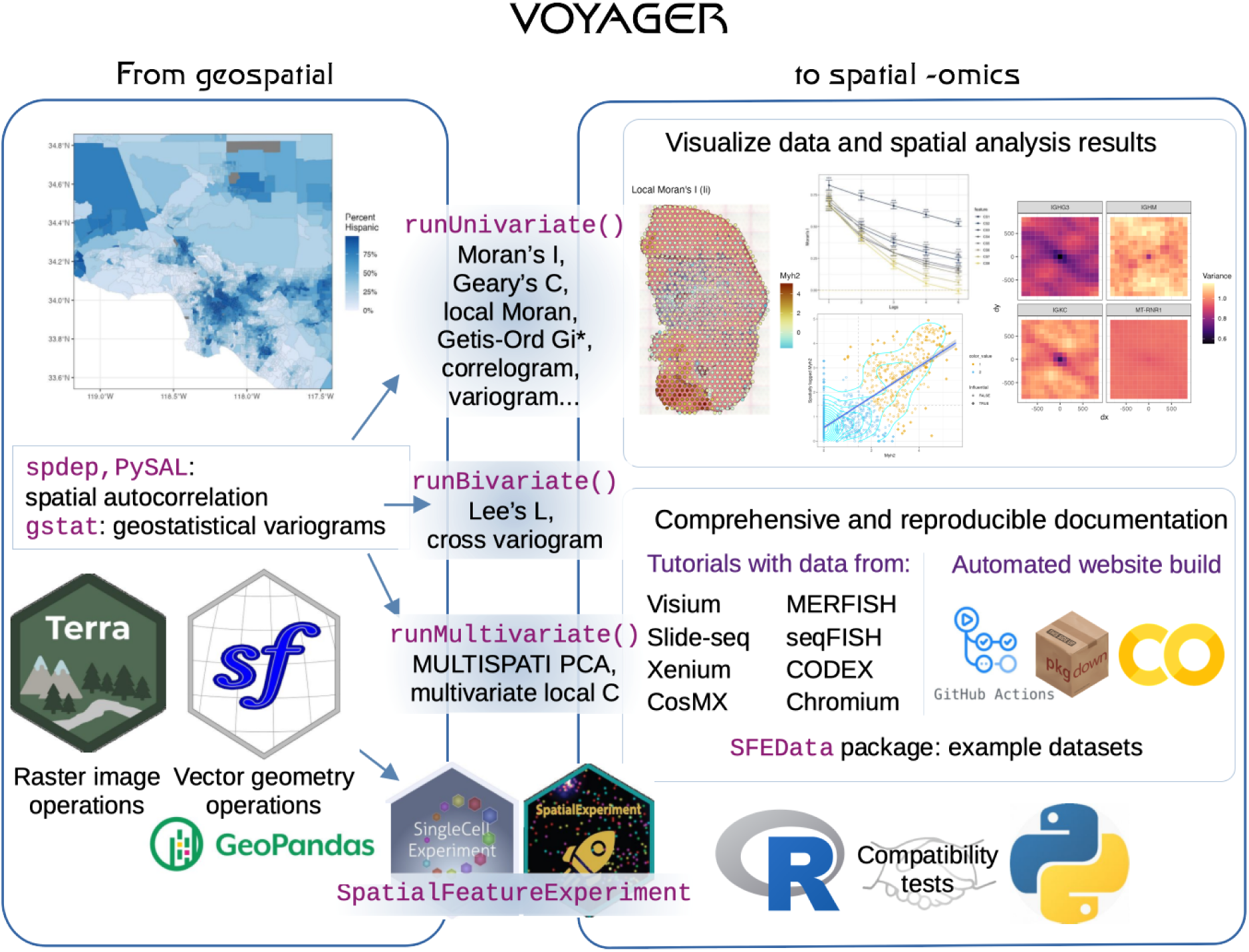
Schematic overview of the Voyager framework. Voyager brings exploratory spatial data analysis (ESDA) methods initially developed for geospatial data to spatial -omics, with a consistent user interface for different methods. Voyager is based on the SpatialFeatureExperiment (SFE) object. In R, SFE uses sf and terra to extend SingleCellExperiment (SCE) and SpatialExperiment (SPE). In Python, SFE extends AnnData with GeoPandas. Voyager implements plotting functions for gene expression, cell attributes, and spatial analysis results. Spatial results shown in this schematic are local Moran’s I (left), correlogram (center top), Moran scatter plot (center bottom), and variogram map (right). The documentation website includes tutorials that demonstrate ESDA on data from multiple spatial -omics technologies, including Visium, Slide-seq, Xenium, CosMX, MERFISH, seqFISH, and CODEX. The website is built automatically with GitHub Actions for reproducibility, and Google Colab notebooks are automatically generated from the vignettes. Compatibility tests are used to make sure that the R and Python implementations return consistent results for core functionalities.

Spatial data analysis methods can be categorized as neighborhood or distance view; the former uses a spatial neighborhood graph to indicate spatial adjacency, while the latter uses physical distance^12^. Voyager wraps many spatial methods from widely used packages, such as spdep (neighborhood) and gstat (distance) in R, and PySAL in Python. ESDA methods can also be broadly categorized by the number of variables analyzed: univariate (e.g. Moran’s I^16^), bivariate (e.g. Lee’s L^25^), and multivariate (e.g. MULTISPATI PCA^26^). In Voyager, each of these has a main function providing a uniform user interface to a variety of methods, thereby simplifying access for users (Figure 1). This structure was inspired by the Tidymodels machine learning framework^40^. These methods can also be categorized as global, where one set of results is returned for the entire dataset, or local, where each location or cell has its own set of results.

The latter facilitates explorations of local heterogeneity in spatial relations. Genes can have different length scales of spatial autocorrelation, which can be explored with the correlogram^21^ and variogram^22^ (Figure 1). The length scale can differ in different directions, i.e. exhibit anisotropy, which can be explored with anisotropic variograms and variogram maps^22^ (Figure 1). Users can extend Voyager and make the uniform user interface run custom ESDA methods to reduce redundant code and facilitate organization and visualization of results. Hypothesis testing is implemented for some of these methods to identify genes or regions whose spatial characteristics are unlikely to occur by chance (Table 1). However, since the assumptions behind some of the tests might not hold well for non-normal gene expression data, significant results should be interpreted as indicating “interesting” genes or regions.

Geospatial data tend to have a much smaller number of features and observations than modern single-cell spatial -omics datasets, so Voyager implements parallel processing when running a univariate or bivariate spatial method over a large number of genes. Voyager also reimplements methods whose current implementations don’t scale to modern spatial -omics data, thereby drastically speeding up computation. These methods include MULTISPATI PCA, Lee’s L, finding bounds of Moran’s I from spatial neighborhood graphs^20^, and distance-based edge weighting of k-nearest-neighbors or distance-based graphs (Supplementary Figures 1-2).

Visualization is an essential part of the EDA process. Voyager implements static plotting functions for gene expression, cell attributes, and cell projections along dimensions obtained by dimension reduction plotted in histological space. The histology image can be optionally plotted behind the cells, or in the case of Visium technology, the spots. While most existing packages plot cells as points, Voyager can plot cell or nuclei segmentation polygons as well. For larger datasets, users can specify a bounding box to zoom into a smaller area. The default palettes are designed to have color values discernible with color vision deficiencies (Supplementary Figure 3). Default continuous palettes come from ColorBrewer^41^, scico^42^, and viridis. A sequential palette is used by default, but a divergent palette is available when there is a meaningful center of divergence, such as 0 in local Moran’s I and Lee’s L. The default categorical palette comes from dittoSeq^43^, which was designed for colorblind-friendly scRNA-seq data visualization. In addition, Voyager implements plotting functions for spatial analysis results, such as the Moran scatter plot, correlograms, variograms, and local spatial statistics shown in histological space (Figure 1).

Voyager has a comprehensive documentation website that features tutorials on applying EDA and ESDA to datasets from multiple spatial -omics technologies, including 10X Visium and Xenium^44^, Nanostring CosMX^45^, Vizgen MERFISH^6^, Slide-seq^46^, seqFISH^47^, and CODEX^48^ (Figure 1). On the website, each -omics technology has a landing page with an introduction to the technology and a table linking to tutorials using a dataset from the technology. Each ESDA method has a similar landing page, with an introduction to the method and a similar table, linking to sections in each tutorial using the ESDA method, some of which include further considerations on the ESDA methods and references to the geospatial ESDA literature. Example datasets used in the tutorials are available as SFE objects in the SFEData package. In addition, there are tutorials instructing users on constructing an SFE object and extending Voyager for custom ESDA methods.

While Voyager is focused on spatial data, neighborhood view ESDA methods can be applied to the k-nearest-neighbor graph in gene expression PCA space. This is illustrated in depth for a 10X Chromium and a Visium dataset. Basic tutorials also introduce analysis for Split-seq, ATAC-seq, 10X single cell multiome ATAC + gene expression, ClickTags, 10X single nuclei, and 10X single cell CRISPR screen performing data preprocessing and quality control.

To ensure that the tutorials are reproducible, Voyager builds the documentation website on GitHub Actions, rendering all the tutorials on the cloud. Because the GitHub Actions runner has fewer computational resources than a typical modern laptop, this ensures that the vignettes are scalable to larger datasets, such as a MERFISH mouse liver dataset with almost 400,000 cells. The tutorials are also automatically converted into Google Colab notebooks to be run interactively, allowing users to experiment with different parameters, and facilitating exploration of new datasets and customization of workflows.

In order to ensure that analysis results are not language dependent, as is the case currently in single-cell and spatial -omics data analysis where results differ depending on whether analysis was performed in R or Python, we have developed a Python implementation of core functionalities of the more comprehensive R package Voyager. The implementation, called VoyagerPy, is equivalent to the R implementation, as ensured via compatibility tests. Because Bioconductor requires software packages to have unit tests and pass a daily automated check, whereas PyPI and conda do not perform automated tests, the Python implementation is thus indirectly held to the Bioconductor standard. VoyagerPy has the added advantage of facilitating the utilization of deep learning and image analysis methods for spatial -omics, as there is better support for such methods in Python than R. The Python implementation also supports visualization in Matplotlib.

### ESDA case studies

Select sase studies taken from the documentation website showcase the potential biological insights that can be gleaned from ESDA. First, we examined a mouse skeletal muscle Visium dataset^49^, two days after notexin injury (Figure 2A-C). In the H&E image, the region with many blue leukocyte nuclei is the injury site, the darker red strip and blocks are muscle-tendon junctions, and the remaining pink regions are myofibers (Figure 2A). When the library size (nCounts) per spot among spots that intersect the tissue is plotted in space, we find that different histological regions have different library sizes. For example, the muscle-tendon junctions tend to have smaller library sizes than myofibers and part of the injury site, and regions with tightly packed myofibers (top and bottom left) tend to have larger library sizes than the region with larger myofibers surrounded by leukocytes (right). While there is no one-to-one correspondence between Visium spots and myofibers, geometric operations can find the myofibers that intersect each Visium spot and their areas using myofiber segmentation. In this dataset, the library size and number of genes detected (nGenes), often used as QC metrics, are related to myofiber size – spots on larger myofibers tend to have more genes detected given the same library size than spots on smaller myofibers (Figure 2A-B).

**Figure 2:**
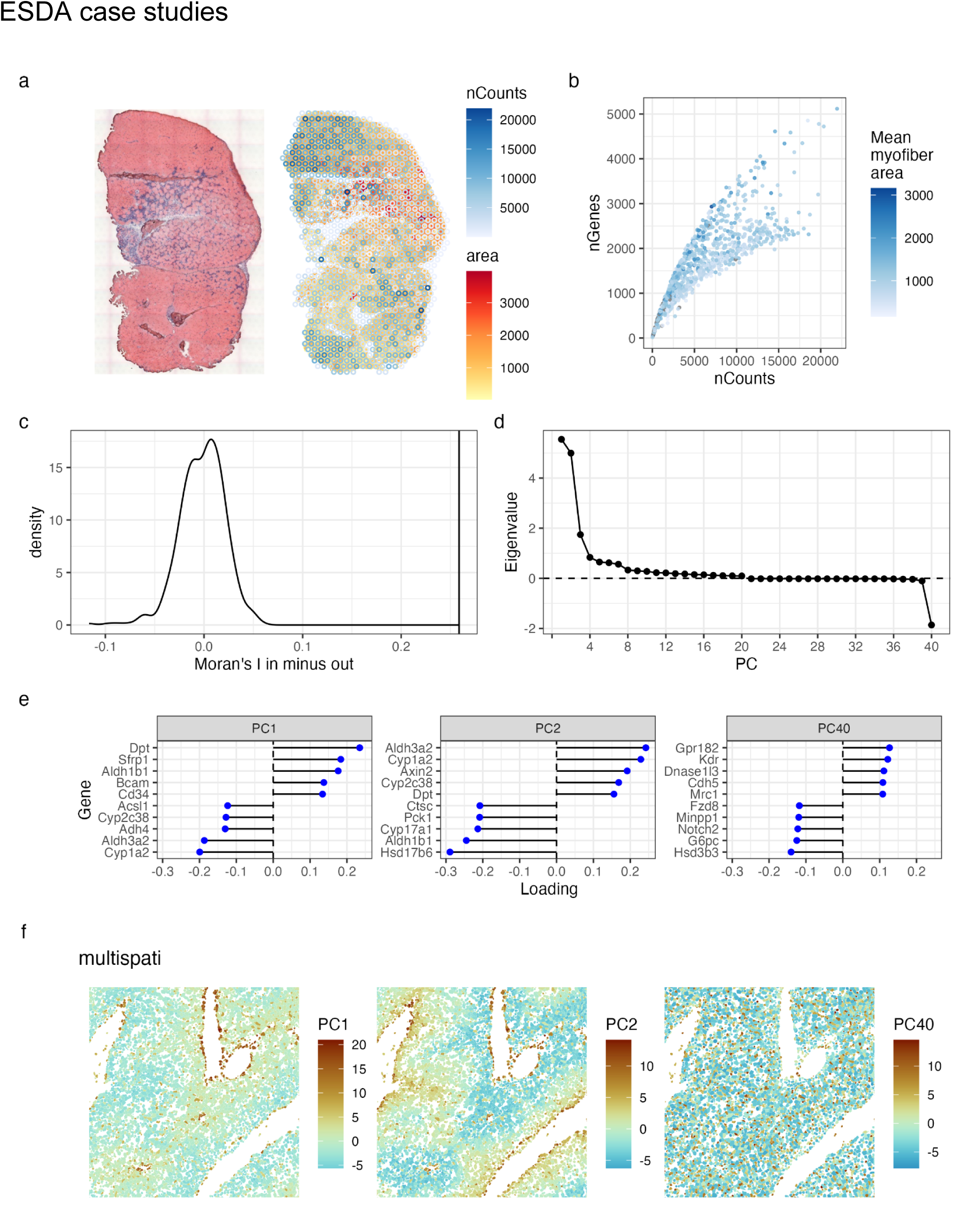
Applications of Voyager on spatial transcriptomics datasets. A) In a mouse skeletal muscle dataset, the total UMI counts, or library size per spot (nCounts), are plotted in space as blue open circles and myofibers are colored in red according to their cross section areas. Only spots that intersect tissue are plotted. The H&E image is plotted on the side as a reference. B) Scatter plot of the number of genes detected per spot (nGenes) vs. nCounts, colored by mean area of myofibers that intersect each spot. C) Simulated (density plot) and observed (vertical line) difference between Moran’s I in nCounts of spots that intersect tissue (in) and that of spots that don’t (out). D) The 20 most positive and 20 most negative eigenvalues from MULTISPATI PCA of a mouse liver MERFISH dataset. As other eigenvalues were not computed, there is a break after PC20 in this plot. E) The most positive and negative gene loadings for PCs 1, 2, and 40. F) A subset of the MERFISH data showing a portal triad (near top right) and two central veins (left and bottom right), with cell polygons colored by their projections into 2 PCs with the most positive eigenvalues and the PC with the most negative eigenvalue (“PC40”). The first 2 PCs show zonation.

Moreover, the library size in the tissue has stronger spatial autocorrelation than outside the tissue, as indicated by a larger positive Moran’s I (Figure 2C, Supplementary Note 2). The library size values are permuted in space for spots that intersect the tissue and those that don’t, and Moran’s I is computed for these permutations to estimate a null distribution. The density plot in Figure 2C shows the null distribution of permuted Moran’s I from spots intersecting tissue minus that from spots not intersecting tissue. The vertical line indicates the actual difference, which is much larger than all 499 simulated values. This confirms the finding that in spatial transcriptomics, library size is biologically relevant and should not be treated as a technical artifact as is commonly done for scRNA-seq ^50^.

The ESDA method MULTISPATI PCA is particularly interesting, as we see from analysis of a mouse liver MERFISH dataset from the Vizgen website^51^ (Figure 2D-F, Supplementary Note 2). While non-spatial PCA maximizes variance explained by each principal component (PC) given that the PCs are orthogonal, MULTISPATI PCA maximizes the product of variance explained and Moran’s I, which is the eigenvalues (Figure 2D). Positive eigenvalues mean that the PCs not only explain more variance, but also are spatially coherent (large positive Moran’s I).

Negative eigenvalues mean that the PCs not only explain more variance, which is non-negative, but also have negative spatial autocorrelation, i.e. nearby values tend to be more different. In this dataset, the positive eigenvalues show an elbow as in non-spatial PCA, and there is one substantial negative eigenvalue (Figure 2D). In non-spatial PCA, the PCs are not spatially structured until PC5, which picks up zoning (Supplementary Figure 4). PC1 highlights Kupffer cells (Cdh5) and endothelial cells (Egfr), and PC2 also highlights endothelial cells (Supplementary Figure 4A). In contrast, because MULTISPATI PCA also maximizes Moran’s I, zonation is picked up by the first 2 PCs. PC1 is periportal and PC2 is pericentral (Figure 2E-F). This may complement existing methods to find spatially variable genes (maximize Moran’s I) that are also more likely to be biologically relevant (maximize variance explained). Furthermore, spatially coherent PCs can be used for clustering to find more spatially coherent clusters in some cases, complementing clusters found with non-spatial PCs (Supplementary Figures 5-6, Supplementary Note 2).

Negative spatial autocorrelation is one of the most neglected topics in spatial data analysis^52^, as there are many more examples of positive than negative spatial autocorrelation. Negative spatial autocorrelation can arise from competition between neighbors (see^52^) or from functional roles played by spatial contacts between different types of entities. In this dataset, the latter seems to be the case: the PC with the most negative eigenvalue separates endothelial cells (Kdr) and Kupffer cells (Cdh5) from hepatocytes (Hsd3b3, Figure 2E-F). Existing methods of spatially informed dimension reduction^53–55^ and methods to find spatially variable genes^56–59^ tend to only consider positive spatial autocorrelation. This example shows that negative spatial autocorrelation is relevant to biology at the single-cell level and should be further investigated, as the cells, unlike Visium spots and administrative boundaries, are meaningful and non-arbitrary units of observations and biological function.

Finally, we note that neighborhood view spatial methods can be applied to neighborhood graphs in gene expression space rather than histological or geographical space. We applied spatial statistics to quality control metrics and gene expression in a 10X Genomics Chromium peripheral blood mononuclear cells (PBMC) dataset^60^ using the k-nearest-neighbors graph in PCA gene expression space as the “spatial” neighborhood graph (Figure 3). Moran scatter plot of library size shows further evidence that library size is confounded by biology even in non-spatial data (Figure 3B). In a Moran scatter plot, the x-axis is the value of a variable at each cell, and the y-axis is the spatially lagged value (i.e. sum of values from spatial neighbors weighted by edge weights of the spatial neighborhood graph). When the adjacency matrix of the spatial neighborhood graph is row normalized, it is shown in Anselin 1996^12^ that the slope of the line fitted to the scatter plot is global Moran’s I, while the scatter plot shows local heterogeneity in spatial autocorrelation. Here, above the fitted line, cluster 5 (activated T cells) tends to have larger library sizes and stronger “spatial” autocorrelation in library size than average (Figure 3B).

**Figure 3:**
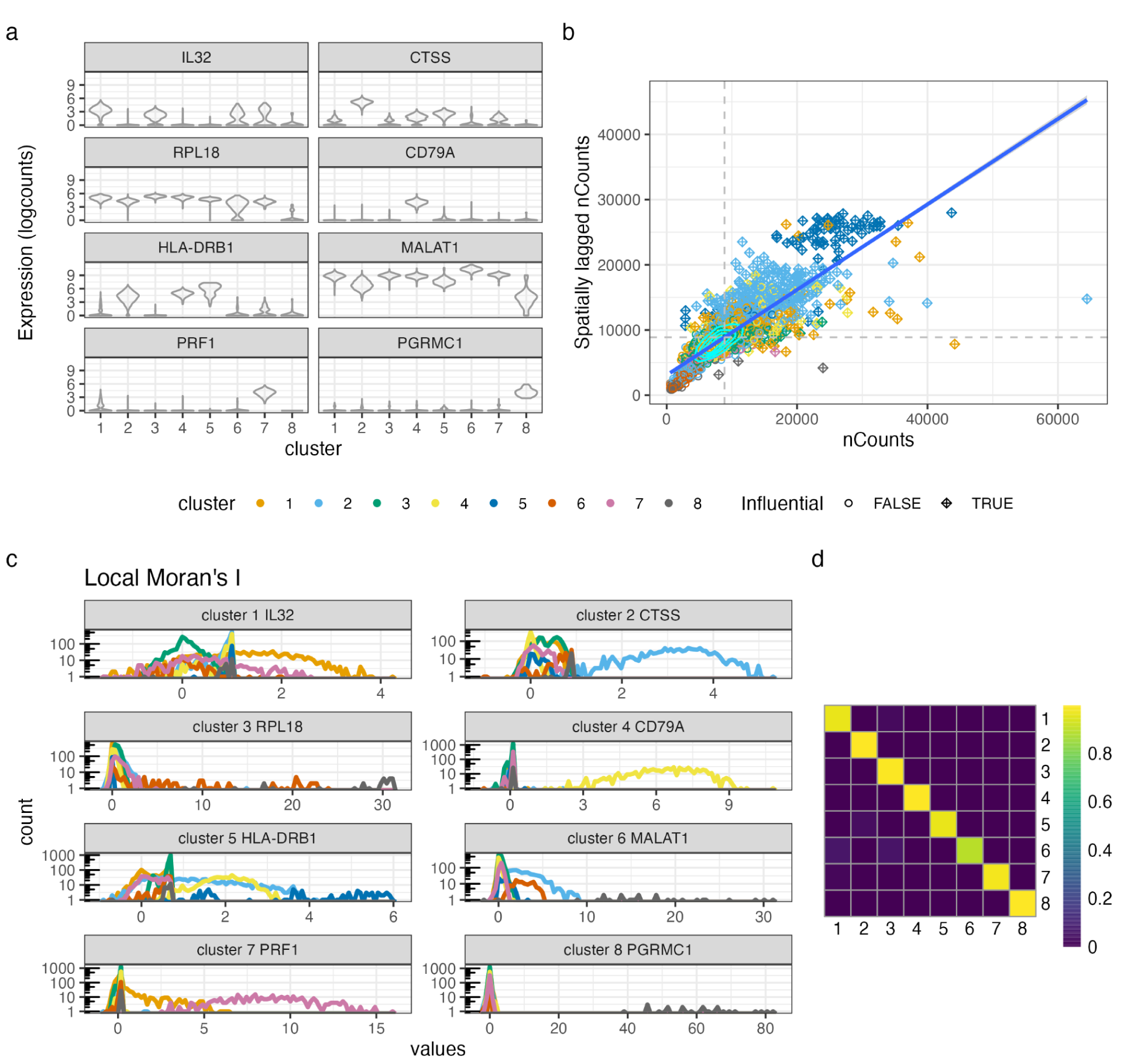
Application of neighborhood view spatial statistics on non-spatial scRNA-seq. A) Violin plots of log normalized counts of the top marker gene of each Leiden cluster in the PBMC dataset. B) Moran scatter plot of nCounts in a 10X Chromium human PBMC dataset. The spatial lags were computed with the k nearest neighbors graph in PCA gene expression space. Colors indicate clusters, and point shape indicates whether the point is influential to the fit of the blue line, which is the least square fit to the scatter plot. The gray shade around the line is the 95% confidence interval of the fit. Contours show the area with the highest point density. The gray dotted lines show the mean on the x and y axes. C) Histograms of local Moran’s I values per cell of top marker genes of each cluster in the PBMC dataset, colored by cell cluster. The y axis (number of cells per bin) is log-transformed for better dynamic range. The histograms are plotted as lines instead of bars to avoid overlapping bars from different clusters. D) Concordex heatmap for the PBMC Leiden clusters. High diagonal and low off diagonal values indicate high clustering quality, or that the Leiden clusters reflect the k-nearest-neighbor graph well, but cluster 6 has somewhat lower quality.

Local Moran’s I is a locally disaggregated form of Moran’s I, that measures the contribution of each cell to Moran’s I^23^ (Supplementary Note 2). Positive values indicate neighborhoods homogeneous in the variable of interest, and negative values indicate heterogeneous neighborhoods. We computed local Moran’s I (Ii) for the top marker gene of each cluster (Methods). The marker gene has much higher Ii in cells in the cluster of interest than cells in other clusters, except for clusters 3 and 6, whose top marker genes don’t clearly distinguish these clusters from most other clusters and don’t seem to have cell type-specific functions (Figure 3A, C). When the marker gene is highly specific, cells in other clusters display an Ii tightly clustered around 0, as shown for clusters 4 (B cells) and 8 (platelets). When the marker gene is not very specific, cells in other clusters that also express the marker gene often display an Ii higher than clusters not expressing the gene (e.g. cluster 1 T cells and cluster 7 natural killer cells and cytotoxic T cells for IL32 and PRF1; cluster 5 activated T cells, cluster 2 monocytes, and cluster 4 B cells for HLA-DRB1, Figure 3A, C).

However, Ii as an index of local homogeneity, does not always correspond to expression level, hence revealing additional nuances of the clusters and marker genes. For example, CTSS, a marker gene of cluster 2 which is also somewhat expressed in clusters 4 and 5, is not more homogeneous in clusters 4 and 5 than clusters with lower expression (Figure 3A, C). Furthermore, cluster 1 marker IL32 has locally negative Ii among some cells in clusters 1, 3, 6, and 7 that have higher expression, indicating local heterogeneity, while unlike for the other marker genes, cells in clusters with low expression of this IL32 have somewhat positive Ii, indicating homogeneity (Figure 3C). The Concordex metric was devised to quantify k-nearest-neighbor graph based cluster quality as an alternative to UMAP, by indicating how well the clusters match the graph structure ^61^. As Leiden clustering is graph-based, Ii on cluster marker genes can give nuances to the single numbers from Concordex and to the quality of the marker genes (Figure 3D). While Ii has been used to identify spatially variable genes^62^, this case study shows that it can potentially be applied to the k-nearest-neighbor graph in gene expression PCA space as another form of differential expression (DE). While these univariate methods can only be applied to one out of thousands of genes at a time, they can give a more nuanced view of genes of interest discovered by other methods as shown here, or applied to features derived from multiple genes such as dimension reduction cell or spot embeddings to explore spatial characteristics of gene programs (Supplementary Figures 5-6).

### Compatibility tests

In scRNA-seq, Seurat and scanpy are both commonly used EDA frameworks. However, their default settings not only yield different log fold changes^63^ but also highly divergent PCA results, as exemplified using a mouse olfactory bulb Visium dataset^64^ (Figure 4A, C-D), which might lead to different biological conclusions. To avoid such discrepancies, Voyager implements extensive compatibility tests to ensure that the R and Python implementations return the same results for core functionalities. Thus, there are no visible differences in Visium spot embeddings in the first two PCs returned by the scater (Voyager R workflow) and VoyagerPy implementations (Figure 4B). In Seurat vs. scanpy, the cosine difference (see Methods) between PCA eigenvectors (gene loadings) in each of the top 50 PCs is nearly 1 after the first few PCs, which means the angles between the corresponding eigenvectors are nearly 90 degrees (Figure 4C). In Voyager, this difference is much smaller, well below machine epsilon (dashed line, around 1.5e-8, see Methods) until PC20 (Figure 4C). While the difference sometimes rises above machine epsilon after PC20, it does not exceed 1e-5. While Seurat and scanpy implementations of PCA with default parameters produce sizable differences in the proportion of variance explained by each PC, these differences in the R and Python implementations of Voyager are generally within machine epsilon (Figure 4D).

**Figure 4:**
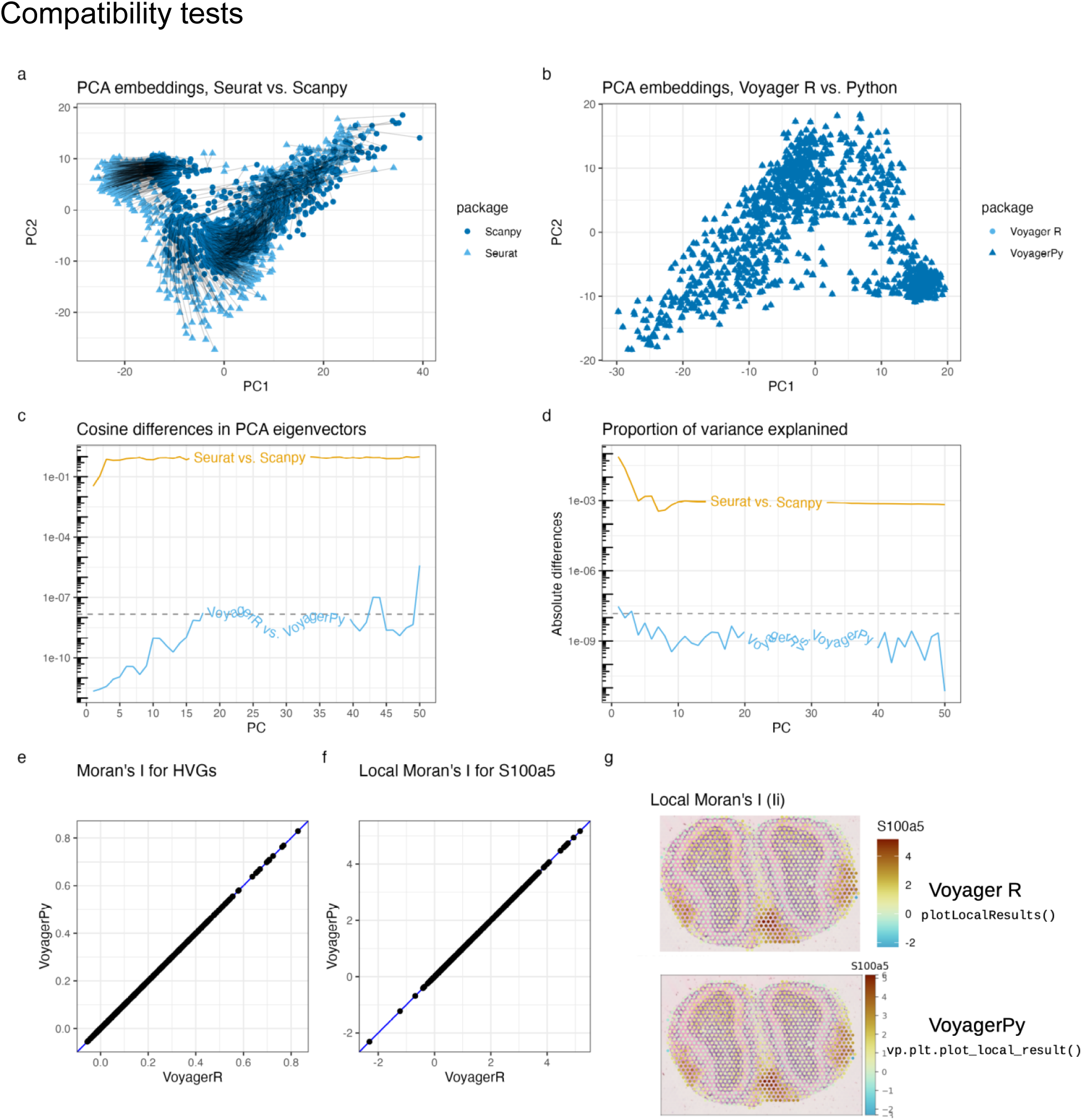
Comparisons between results obtained by Seurat and scanpy, and between Voyager R and Python for a mouse olfactory bulb Visium dataset. A) Comparison of Visium spot embeddings in the first 2 PCs from Seurat and scanpy with default parameters. The lines connect corresponding spots in Seurat and scanpy. B) As in A, but for Voyager R and VoyagerPy, with parameters stated in this section. C) Cosine distances between the first 20 PCA eigenvectors (gene loadings) from Seurat and scanpy (yellow), and from Voyager R and Python (blue). The dashed line is the magnitude that can be explained by machine double precision. The text part of the line is somewhat smoothed for readability but should not affect interpretation. D) Absolute values of differences in the proportion of variance explained by each of the top 20 PCs. E) Moran’s I from VoyagerPy vs. Voyager R. The blue line is y = x, showing that the results are consistent. F) Same as E but for local Moran’s I for gene S100a5. G) Plotting the local Moran’s I values in space, with the H&E image behind the spots, from Voyager R (top) and VoyagerPy (bottom).

For spatial statistics, the R and Python implementations of Voyager also produce consistent results for global Moran’s I of the highly variable genes (Figure 4E), with differences within epsilon (1.61e-17, see Methods). For local Moran’s I (Figure 4F-G), the results are the same (within epsilon, 3.89e-16) but with a non-default spdep parameter (Supplementary Note 3). In both implementations, besides the different default styles of `ggplot2` and `matplotlib`, the plotting functions produce visually similar plots with the same palettes (Figure 4G).

While the R and Python implementations of Voyager may eventually diverge in functionalities, as the two languages have better support for different types of analyses, the compatibility tests will continue to ensure that the core functionalities and basic vignettes return the same results in R and Python, so language preference does not inadvertently lead to different interpretations of data.

Notably, Seurat and scanpy produce different PCA results largely due to their different methods of finding highly variable genes. However, the PCA results remain somewhat different even when using the same set of highly variable genes because of a hidden default in Seurat that clips scaled data to 10, while scanpy by default does not clip the scaled data (Supplementary Figure 7). Moreover, Seurat and scanpy return different marker gene rankings from ostensibly the same DE method (e.g. t-test or Wilcoxon test) because they rank the genes differently and even estimate different log fold changes differently^63^ (Supplementary Note 4). Most users may be unaware of such inconsistencies and the hidden parameter defaults and method choices that cause them. In the interests of transparency, we have therefore documented Voyager’s default parameters and the reasons for choosing them (Supplementary Note 3). We have also elucidated reasons for the divergent log fold changes in Seurat vs. scanpy and compared effect sizes and p-values from DE with Wilcoxon rank sum test in Seurat vs. scanpy and in scran (Voyager R workflow) vs. VoyagerPy. In contrast to Seurat and scanpy, VoyagerPy and scran give largely consistent results (Supplementary Note 4, Supplementary Figures 8-10).

At present, Voyager implements compatibility testing for two tutorials: a mouse olfactory bulb Visium dataset from the 10X website, and a univariate spatial statistics analysis of the k-nearest neighbor-graph of a human peripheral blood mononuclear cells (PBMC) 10X Genomics Chromium dataset. The defaults used in these tutorials covered by the compatibility tests are listed in Supplementary Note 3; some of the defaults come from conventions and defaults in established packages and hence are subject to further research and improvement.

## Discussion

Software packages for data visualization, general EDA frameworks, and more specialized tasks have proliferated in the field of spatial -omics, but much of the ESDA tradition has not been utilized. As an EDA framework, Voyager is similar to software packages such as Seurat, squidpy, Giotto, and semla, but Voyager is unique in systematically facilitating the porting of decades of ESDA research to spatial -omics, with a consistent user interface. Just as the discovery of overdispersion in RNA-seq data led to the widespread adoption of negative binomial models in transcriptomics^65–67^, better characterization of spatial properties of gene expression in different tissues with ESDA—such as by taking local heterogeneity, length scales, and anisotropy of spatial autocorrelation, and negative spatial autocorrelation into account—can lead to better specialized downstream models such as those identifying spatially variable genes or simulating data for benchmarking^68^.

By building upon the SCE and AnnData infrastructures and ecosystems, Voyager complements many other spatial and non-spatial data analysis methods. The SFE class extends SCE and AnnData with efficient tools from the geospatial field to represent and operate on vector geometries and raster images. Voyager has a comprehensive, reproducible, and easy-to-navigate documentation website with tutorials on data from various technologies and ESDA methods, with references for further reading and considerations from the ESDA tradition. Extensive compatibility tests ensure that the R and Python implementations of Voyager return consistent results for core functionalities and transparency on defaults. The Voyager project also bridges the R vs. Python divide in single-cell and spatial -omics bioinformatics, where hidden defaults and undocumented divergent implementations cause language preference to inadvertently lead to different results that may affect biological conclusions. Thus, even for non-spatial EDA, it provides a substantial advantage over current packages.

The scholar Jaroslav Pelikan wrote that “Tradition is the living faith of the dead, traditionalism is the dead faith of the living.”^69^ While the ESDA tradition was largely developed prior to the rise of spatial -omics, it can help us gain insights by taking the spatial aspects of biological data into consideration. As the ESDA tradition is ever evolving, Voyager is designed to be extensible by users and developers to facilitate the application of novel methods with a consistent user interface. Future versions of Voyager can also take into account the peculiarities of spatial -omics data, such as larger dataset sizes, case and control comparisons, multiple biological replica, non-normal distribution of the data, and 3-dimensional data from thick slices and multiple sections. At present, unlike squidpy, Giotto, and semla, Voyager does not implement ESDA for categorical data (Supplementary Table 1), as this is less developed in the geospatial field^21, 70^. Furthermore, categorical spatial methods using SCE such as lisaClust^71^ can be easily applied without being incorporated into Voyager. However, with more considerations on the nature of cell types^72, 73^ and ESDA of categorical variables, categorical methods may be added in future versions.

Additional future directions include storing geometries and spatial results on-disk. While the SCE infrastructure already allows for on-disk gene count matrices with DelayedArray, the geometries and spatial analysis results are currently stored in memory. Moreover, while we extensively documented the Voyager defaults to avoid inconsistencies between the R and Python implementations, the reasoning behind them is often based on convention in the field. Further research should scrutinize the effects of changing these parameters, such as the type of spatial neighborhood graph and edge weights. The problem of choosing a spatial neighborhood graph has long been studied, and some methods to find a graph based on the data have been devised^74^, but they may or may not be suitable for spatial -omics data. Finally, while we have chosen colorblind-friendly default palettes to make Voyager more accessible, future research should be conducted on the accessibility of spatial -omics data analysis, such as in data sonification.

## Methods

All R plots in the figures were made with R 4.3.0 with Apple vecLib BLAS, Bioconductor 3.17, Voyager 1.2.4, SpatialFeatureExperiment 1.2.1, scater 1.28.0, spdep 1.2.8, Seurat 4.3.0, sf 1.0.12, and ggplot2 3.4.2, on MacOS Ventura 13.3.1, 2.3 GHz Dual-Core Intel Core i5, 8 GB RAM. R package profvis 0.3.7 was used to profile time and memory usage by lines of code in the benchmarks, and bench 1.1.2 was used for the benchmarks over different numbers of cells. When comparing Seurat vs. scanpy and the R and Python implementations of Voyager, the Python code was run through reticulate (v1.28) in RStudio. Python 3.10, scanpy 1.9.3, and VoyagerPy 0.1.1 were used. Multipanel plots were assembled with patchwork 1.1.2 when all panels are R plots, and were otherwise assembled in LibreOffice Draw.

### Spatial methods

At present, all neighborhood view spatial methods are implemented in spdep and wrapped by Voyager, except for Lee’s L, which has a more efficient implementation in Voyager. Defaults follow those of spdep. All distance view spatial methods are implemented in gstat and wrapped by Voyager. Variogram model fitting is implemented in automap, which is a user-friendly wrapper of gstat that tries a number of different models and selects the one with the best fit. Voyager features its own, more efficient implementation of MULTISPATI PCA, and hence does not depend on adespatial which originally implemented MULTISPATI PCA.

In contrast to the plotting functions in spdep and gstat, Voyager plotting functions are based on ggplot2^75^ to be more visually appealing and customizable by users and are designed to visualize results from multiple features and tissue sections at once.

### Compatibility tests

Functionalities such as finding highly variable genes, PCA, and DE have been reimplemented in VoyagerPy to match the implementations in scater and scran used in the Voyager R workflow to ensure that the two implementations of Voyager give consistent results for these procedures.

Everything in the two core tutorials other than the plots themselves—i.e. procedures that yield numeric output, such as PCA and Moran’s I—is subject to compatibility tests to verify that the R and Python implementations of Voyager produce the same results for core functionalities. The plots cannot be quantitatively and automatically compared because of the different default styles and mechanisms of ggplot2 and matplotlib, so the comparison is performed based on visual similarity. The “epsilon”, or numeric differences that can be accounted for by machine double precision, was established as `sqrt(.Machine$double.eps)` in R. To compare PCA eigenvectors (gene loadings), cosine difference is used to geometrically compare the vectors. This is measured as the magnitude of difference between the cosine of the angle between the two vectors and 1, i.e. cosine of 0 and 180 degrees. The value 180 degrees was chosen because the eigenvectors can be flipped and yet produce equivalent results in PCA. This comparison was performed on each of the first 50 PCs individually for Figure 4. To compare the proportion of variance explained, the absolute value of the difference was used.

For the comparisons in Supplementary Note 4 and Supplementary Figures 8-10, the same PBMC 5k dataset used in Figure 3 was used. Standard Seurat and scanpy log normalization was performed. Seurat was used to find HVGs, scale the data so each gene has mean 0 and variance 1, and perform PCA on the scaled data with the HVGs. Fifty PCs were computed, and all were used to find a k-nearest-neighbor graph with k = 20, the default in FindNeighbors(). Then Seurat was used to cluster the cells, with the Louvain algorithm, with resolution = 0.5. Wilcoxon rank sum test was performed with Seurat, scanpy, scran, and VoyagerPy. Default parameters as of versions mentioned in the first paragraph of the Methods section were used, except that for scran, non-default pval.type = “all” and direction = “up” in the findMarkers() function were used, because the only DE functionality implemented in VoyagerPy is equivalent to using these parameters in scran.

### Website build

The R Voyager documentation website is built with pkgdown on GitHub Actions, which builds function references and vignettes from the R package source code. All imported and suggested packages are installed on a fresh machine on the cloud and all vignettes are run on the cloud to be rendered, to ensure that they are reproducible. The Google Colab notebooks are automatically generated from the R Markdown vignettes with another GitHub Action. Because Bioconductor limits the installed size of the package, which includes the rendered vignettes, the vignettes on the documentation website are in a documentation branch separate from the main and devel branches that sync with Bioconductor, while a shorter vignette is on Bioconductor. Also, there are packages suggested in the documentation branch but not the main branch, as while they are used in vignettes on the website, they are not used in the Bioconductor vignette or the package itself. The code in the documentation branch is synchronized with code from the main branch by merging from the main branch, but the documentation branch is never merged into the main branch.

The VoyagerPy website is built with sphinx and deployed via GitHub Actions.

### Performance improvements

In the benchmarks, a mouse liver MERFISH dataset from the Vizgen website with over 390,000 cells after QC was used. After removing cells with a high proportion of transcripts from blank barcodes (and removing the blank barcodes themselves), the dataset was subsetted with bounding boxes of different sizes to produce datasets of different sizes while preserving spatial relationships among cells, which were used in the benchmarks.

### K-nearest-neighbors with inverse distance weighting

The spdep implementation of distance-based edge weights is slow because while spdep uses an efficient implementation of k nearest neighbors and distance neighbors in dbscan, it discards the distances between neighbors returned by dbscan. As a result, spdep has to re-compute the distances to compute the edge weights, which is time consuming (Supplementary Figure 2A). The implementation in SFE uses BiocNeighbors to find the k nearest and distance-based neighbors, allowing users to choose from a number of different algorithms. Then the distances are saved for edge weight computations, skipping the most time-consuming step. While dbscan is not much slower than BiocNeighbors when finding the neighbors (Supplementary Figure 2A-B), we found that not recomputing the distances speeds up finding the spatial neighborhood graph from 8 to over 30 folds and uses over 25 times less memory (Supplementary Figure 1A-B).

### MULTISPATI PCA

The adespatial implementation of MULTISPATI PCA uses base R eigen decomposition, which always computes all eigenvalues and eigenvectors. Subsequently, when the user specifies a small number of eigenvectors, adespatial discards the remaining eigenvectors. Furthermore, adespatial in fact performs the eigendecomposition twice. The first time is in dudi.pca, which performs non-spatial PCA, whose results are passed to the multispati function, which takes some weights and the original data but not the eigenvalues or eigenvectors from the dudi.pca output, and then performs eigen decomposition of the spatially weighted covariance matrix (Supplementary Figure 2C). The implementation in Voyager uses RSpectra for partial eigendecomposition of the spatially weighted covariance matrix, only for the eigenvectors the user requested and only once, hence avoiding a lot of unnecessary computation, speeding up computation by 2 to 20 folds, and using over 5 times less memory (Supplementary Figure 1C-D).

### Lee’s L

The spdep implementation of Lee’s L computes both local and global Lee’s L for one pair of genes at a time. As spatial transcriptomics data has hundreds (smFISH based) to thousands of genes (sequencing based), Voyager’s implementation uses matrix operations to make it more efficient to compute Lee’s L for a large number of genes than iterating through each pair. This speeds up computation over 800 fold and uses 100 times less memory, where one thread was used to iterate through all pairs of genes in the dataset for the spdep implementation (Supplementary Figure 1E-F). The inefficiency of spdep’s implementation is not only due to iterating through the pairwise combinations, but also because of a less efficient computation of the spatially lagged values and the sum of edge weights (Supplementary Figure 2E). Iterating through the pairs with the spdep implementation is so slow that it was only run for the smallest subset with 105 cells in the benchmark.

### Moran bounds

Bounds of Moran’s I given a spatial neighborhood graph can be computed from the largest and smallest eigenvalues of the double-centered adjacency matrix of the graph^20^. In the adespatial implementation of the function finding Moran bounds, all eigenvalues are computed. In Voyager’s implementation, RSpectra is used to find only the largest and smallest eigenvalues of the matrix without computing eigenvectors or the other irrelevant eigenvalues, speeding up computation by 4 folds and using 4 times less memory (Supplementary Figure 1G-H). While much of the time was spent on the eigendecomposition in the adespatial implementation, most of the time was spent on double centering in the Voyager implementation (Supplementary Figure 2G-H). Due to double centering, a dense matrix with as many columns and rows as the number of cells is produced. Unless this can be avoided, this computation consumes a lot of memory for larger datasets. As a result, neither implementation scaled beyond around 6000 cells.

### ESDA case studies

#### Mouse skeletal muscle Visium data

Space Ranger processed data were downloaded from GEO accession GSE161318, sample Vis5A, 2 days after notexin injury. Myofiber segmentation was performed manually with the LabKit ImageJ plugin, exported as TIFF raster, and converted to polygons with R package terra. Redundant vertices of the polygons were removed when the polygons were simplified with rmapshaper (v0.4.5). Visium spot polygons were found from the centroids and spot diameters in pixels in the full-resolution image from the Loupe image alignment JSON file. The tissue was segmented by thresholding the H&E image and removing small pieces. Then, the thresholded mask was converted into polygon with terra, which was used to find spots intersecting the tissue with sf. The gene count matrix and polygons were made into an SFE object available to download from the SFEData package. The sf package was used to find myofiber areas and to determine which myofibers intersected each Visium spot polygon. Only the full-resolution H&E images were available on GEO; to facilitate reproducibility of the vignette and examples, the image was downsampled to fit into a 1024x1024 pixel box. Moran’s I permutation tests were performed on nCounts for spots intersecting the tissue and spots not intersecting the tissue separately, with 499 permutations. The values are permuted in space. Subsequently, the simulated Moran’s I’s from spots not intersecting tissue were subtracted from those from spots intersecting tissue to form a null distribution of this difference. The “observed” value was the observed Moran’s I from spots not intersecting tissue subtracted from that of spots intersecting tissue. Leiden clustering (Supplementary Figure 5) was performed on the first 30 non-spatial and MULTISPATI PCs, with resolution 1 and objective function “modularity” in the R package bluster v1.10.0. Neighbor purity was computed with the neighborPurity() function in bluster. Concordex was computed with the concordexR package v1.0.0. Neighbor purity and Concordex were computed with the k-nearest-neighbor graph (k=6 not including self because of the hexagonal grid) in histological space, although the graph in PCA space was used to generate the clusters. This way the cluster quality indices displayed spatial coherence of the clusters.

#### Mouse liver MERFISH data

The gene count matrix, cell metadata, and cell segmentation polygons were downloaded from Vizgen’s website, and parsed into an SFE object, which is available in the SFEData package. The scuttle package (1.10.0) was used to remove low-quality cells. The proportion of transcripts attributed to blank barcodes was computed, log2 transformed, and cells with log proportion more than 3 median absolute deviations (MADs) higher than the median were deemed low quality and removed. Subsequently, the filtered gene count matrix was normalized by logNormCounts() in scater, and the genes were scaled and centered before performing non-spatial PCA with IRLBA through scater, and MULTISPATI PCA. MULTISPATI PCA requires a spatial neighborhood graph and a k-nearest-neighbor graph with k = 5 (not including self) was used; see Supplementary Note 3 for reasons behind the parameters chosen. Leiden clustering, neighbor purity, and the Concordex index were computed as in the mouse skeletal muscle Visium dataset, except that the first 20 PCs were used with the same spatial neighborhood graph as in MULTISPATI PCA.

#### Chromium PBMC data

The filtered 5k PBMC NextGem v3 data, processed with Cell Ranger 3.0.2, was downloaded from the 10X Genomics website and loaded into R as an SCE object, and was then converted to SFE for “spatial” analyses. Cells with at least 20% of UMIs from mitochondrially encoded genes were removed. Highly variable genes (HVGs) were found with the scran method but without the Lowess fit. The 2000 genes with the highest biological component were used for PCA. The data was normalized with logNormCounts() in scater, and the genes were scaled and centered before performing PCA with the IRLBA algorithm. Based on the variance explained elbow plot, the 10 PCs were used to build a k nearest neighbor graph, with k = 10 (not including self). For Leiden clustering, k = 10 was also used so the clustering results can be compared to the “spatial” results. The objective function selected was “modularity” and the resolution parameter was 0.5. For the “spatial” neighborhood graph, inverse distance weighting and W style normalization were used, for reasons similar to those in k-nearest-neighbor graphs in histological space explained in Supplementary Note 3. The R package scran was used for DE, with pval.type = “all” and direction = “up” in the findMarkers() function to identify genes more highly expressed in each cluster of interest than in the rest of the cells.

## Data and code availability

The mouse skeletal muscle Visium dataset and the mouse liver MERFISH datasets are available in the SFEData R package. These are the GitHub repositories and websites related to this paper:

Voyager R package: https://github.com/pachterlab/voyager

SpatialFeatureExperiment: https://github.com/pachterlab/SpatialFeatureExperiment

SFEData: https://github.com/pachterlab/SFEData

Voyager Python package: https://github.com/pmelsted/voyagerpy

Voyager R documentation website: https://pachterlab.github.io/voyager/

Voyager Python documentation website: https://pmelsted.github.io/voyagerpy

Compatibility tests: https://github.com/pachterlab/voyager-testing

Code to reproduce figures in this paper: https://github.com/pachterlab/MEJLBAMP_2023

## Acknowledgements

We thank SpatialExperiment author Dario Righelli for suggestions in the early development of SFE, and Alik Huseynov for contributions to SFE. This work was supported in part by the Icelandic Research Fund Project [218111-051] and NIH 5UM1HG012077-02. We thank the Caltech Bioinformatics Resource Center for providing computing resources during the development of the project. We thank Sean Davis, the Institute for Systems Biology Cancer Gateway in the Cloud (ISB-CGC), Alex Mahmoud, NSF Jetstream 2 cloud ACCESS allocation BIR190004, and the Bioc organizing committee for computational platforms to host workshops on Voyager and SFE during the development of the project.

## Author contributions

L.M. conceived the main ideas, implemented the R packages SpatialFeatureExperiment, Voyager, and SFEData, wrote most of the R Voyager tutorials, wrote the spatial methods landing pages on the R website, built the R documentation website, and drafted the manuscript. S.A. implemented the foundation for VoyagerPy. P.H.E. implemented VoyagerPy, built the VoyagerPy documentation website, and implemented compatibility tests for the two core tutorials. K.J. wrote some of the R Voyager tutorials and technology landing pages. L.L. implemented the GitHub actions to automatically generate Google Colab notebooks from the R Markdown tutorials, added gget examples in some tutorials, and edited the manuscript. A.S.B. wrote some of the VoyagerPy tutorials, implemented notebooks demonstrating pre-processing of raw data, and edited the manuscript. P.M. conceived the ideas for VoyagerPy. L.P. conceived the ideas of compatibility tests, applying neighborhood view spatial methods to the k-nearest-neighbor graph in PCA space for non-spatial scRNA-seq data, developed the concepts for the landing pages for data collection technologies and spatial data analysis methods, and edited the manuscript. All authors made suggestions to the manuscript.

