## Supplementary notes for "Voyager: exploratory single-cell genomics data analysis with geospatial statistics"

August 19, 2023

#### Supplementary Note 1: The `SpatialFeatureExperiment` class

Voyager is based on the data structure SFE, which in the R implementation inherits from SPE and SCE (Figure 1). As a result, non-spatial scRNA-seq EDA methods and plotting functions implemented in `scater` [1], an existing package for scRNA-seq data preprocessing, quality control (QC), and EDA, can be utilized within Voyager. In particular, Voyager plotting functions are constructed to be consistent with their counterparts in `scater` in order to maintain a uniform user interface. In the Python implementation, SFE analogously builds on top of `AnnData` so other tools using `AnnData` can be applied. This note focuses on the R implementation.

The SFE data structure is the foundation of Voyager's spatial data analyses. SCE is a data structure designed to represent scRNA-seq data. The raw and normalized gene count matrices are in the assays slot, with genes in rows and cells in columns. Any matrix-like class can be used in the assays, including dense matrices, sparse matrices (e.g. `dgCMatrix` from the `Matrix` package), data frames, and `DelayedArray`, which allows for on-disk operations. Cell metadata is in the `colData` field, and row metadata is in the `rowData` field. In addition, analogous to `colData`, cell embedding matrices from dimension reductions are stored in the `reducedDims` field. PCA loadings are attributes of the PCA cell embedding matrix. SCE implements getters and setters for these fields, and getters and setters for the new fields in SFE for geometries conform to the conventions and styles of SCE getters and setters.

SPE is an existing class that extends SCE for spatial -omics data, adding the `spatialCoords` field for spatial coordinates of cell or spot centroids and imgData to organize images associated with the spatial dataset. The images can be on disk or remote and are thus not loaded into memory unless necessary. When the image is read into memory, it is a matrix of color hex codes. In addition, there's a special column in colData called `sample_id` to distinguish between coordinates from different tissue sections, as different sections can have overlapping numeric values of coordinates. The SPE package also implements functions to mirror and rotate the images. Accompanying SPE is the ggspavis package to visualize gene expression and cell attributes in space.

The geospatial tradition is central to how SFE extends SPE. The sf package is the R interface to Simple Features, a standard way to represent vector geometries in the geospatial field. In `sf`, geometries and their attributes can be stored in a data frame with a special column for the geometries. Furthermore, `sf` supports optimized geometric IO and operations with the GDAL and GEOS C++ libraries, respectively. Using geometric operations, the SFE object can be subsetted or cropped with a geometry.

SFE adds `colGeometries`, analogous to `colData`, which is essentially a collection of `sf` data frames, for geometries associated with columns of the gene count matrix such as cells and Visium spots. A cell can be associated with multiple geometries, such as cell and nucleus segmentation polygons. The efficient geometric operations allow the identification of problematic and likely low-quality cells and nuclei with unusual sizes or multiple pieces in quality control. Moreover, we can find characteristics of cells that may be associated with gene expression, such as morphological metrics and proportion of cell area occupied by the nucleus. In data visualization, the cell segmentation polygons can be plotted instead of centroids, revealing the cell morphology and avoiding overlaps between points in areas with high cell density. The `rowGeometries` field

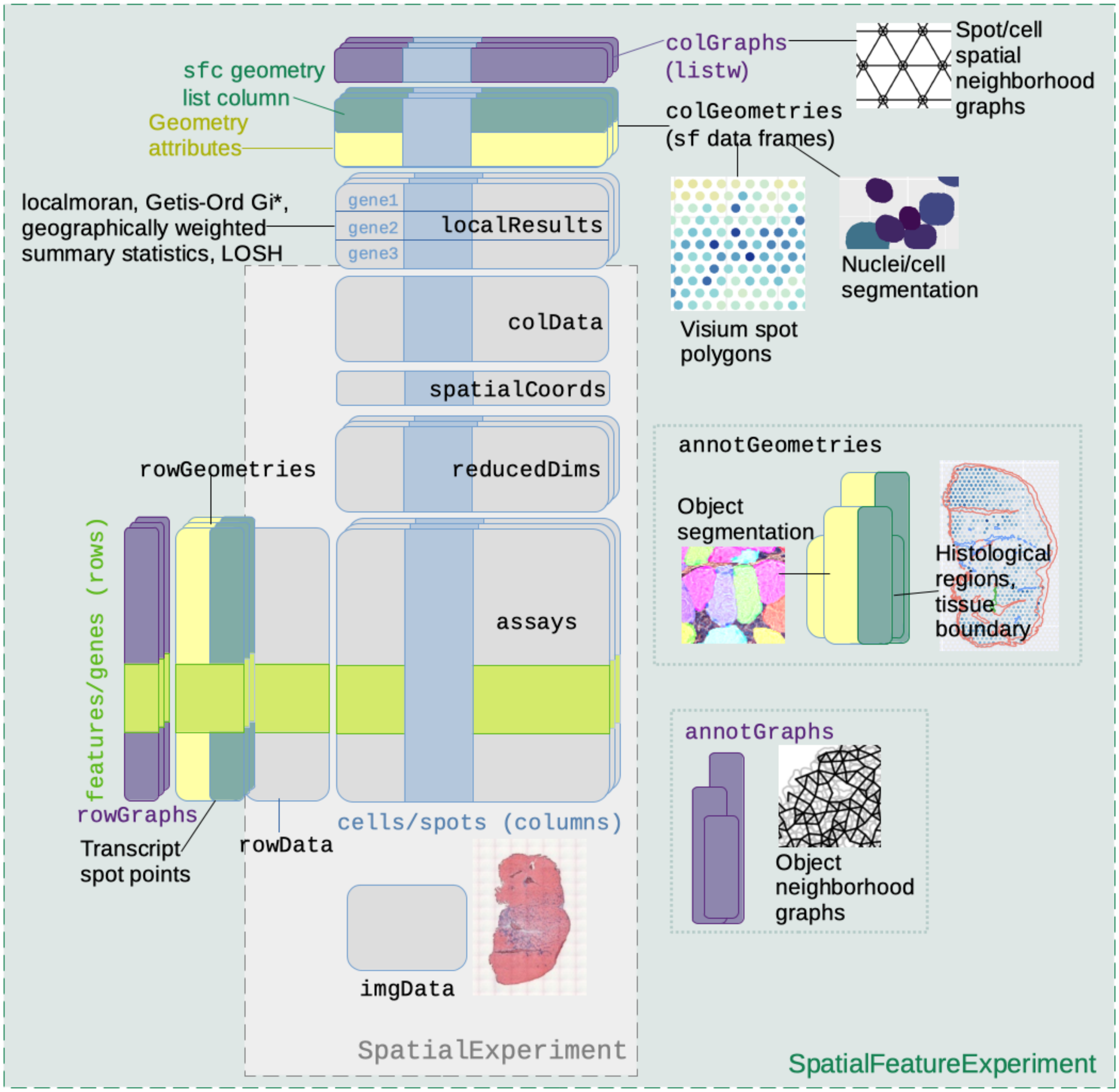


Schematic showing the structure of the `SpatialFeatureExperiment` object and how it extends `SpatialExperiment`.

contains geometries associated with genes or features. It is implemented although not currently used. It can potentially be used for transcript spots from smFISH based datasets, including those not assigned to cells. The transcript spot data is often very large and would benefit from on-disk representations of geometries.

The `annotGeometries` field is for geometries associated with the dataset but does not directly correspond to columns of the gene count matrix. These can be cell segmentation polygons in a Visium dataset, the tissue boundary, or pathologist-annotated histological regions from other software such as QuPath and Fiji/ImageJ. Using geometric operations, we can identify the number of cells or nuclei in each Visium spot, the histological region cells or spots belong to, and the proportion of each Visium spot that is in the tissue or histological region. These can then be related to gene expression in the EDA process. Voyager can compute univariate spatial statistics on numeric columns of `colData`, `colGeometries`, `annotGeometries`, and dimension reduction cell embeddings, in addition to gene expression data.

The neighborhood view of spatial analyses requires a spatial neighborhood graph, whereas SPE does not have a field to organize the graphs. In SFE, the `colGraphs` field stores the spatial neighborhood graph for entities associated with columns of the gene count matrix. The SFE package wraps all methods to find spatial neighborhood graphs in `spdep`, including polygon contiguity, triangulation followed by edge pruning, k-nearest-neighbors, and distance-based neighbors, as well as different types of edge weights, such as binary, row normalization, row and column normalization, and distance based edge weights. In addition, SFE has a much faster implementation of finding distance-based edge weights after finding the k nearest neighbor or distance-based graph. The `annotGraphs` field is for spatial neighborhood graphs of annotation geometries, so spatial analyses can be performed on attributes of these geometries, such as cell area. The `rowGraphs` field is for graphs associated with genes or features, although it is not currently used. The `sample_id` is important here because the spatial neighborhood graph only makes sense within one tissue section. The graphs are represented as `listw` objects as implemented in `spdep`, so no conversion is required when using the numerous `spdep` methods in Voyager.

Results can be organized within the SFE object to link results to features from which they were computed and to facilitate visualization. Local spatial statistics returns one set of results for each cell or spot. To organize the results, the `localResults` field is introduced, analogous to `reducedDims`. Like `reducedDims`, `localResults` are organized by the spatial analysis method, but unlike `reducedDims`, the results for each method are organized by genes or features. Univariate and bivariate local results are stored in the `localResults` field, while multivariate results are stored in `reducedDims` or optionally `colData` if they are vectors or data frames. Univariate global results are stored in `rowData` for gene expression and in the metadata or attributes of the corresponding fields for `colData`, geometries, and dimension reductions. These metadata and attributes can be accessed with getter functions such as `colFeatureData()`. However, bivariate global results are not currently stored in the SFE object due to the great variability in output format.

While the SPE package can read Space Ranger output, SFE does so somewhat differently. Because the Space Ranger output includes spot diameter in pixels in the full-resolution image, SFE constructs Visium spot polygons from the centroids and the diameter. In addition, the pixels can be converted to micrometers, based on the spacing between spots which is known to be 100 $\mu$m. Furthermore, unlike SPE, SFE uses the `terra` package to manage images. The `terra` package is designed for raster geospatial data, which often cannot fit into memory. The images are read as `SpatRaster` objects, which are pointers to the images on disk, so the images are not loaded into memory unless necessary. When the image is plotted, it is not entirely loaded into memory if a lower resolution suffices. When an SPE object is converted into SFE, the images are converted to `SpatRaster`. SFE can also directly read Vizgen MERFISH output, which includes fluorescent images that may not fit into memory, benefiting from `terra`. Furthermore, with `terra`, one can extract values from the raster images with the vector geometries in `colGeometries` and `annotGeometries`.

While SFE objects can be subsetted like a matrix, just like SCE and Seurat objects, because SFE places the geometries front and center, it can be subsetted with a polygon or bounding box, so only data for geometries that intersect the polygon or bounding box are kept. The geometries themselves can also be cropped by the polygon or bounding box.

### Supplementary Note 2: Preliminaries in spatial statistics

For completeness, we briefly review the spatial data analysis methods used in the exploratory spatial data analysis Voyager case studies.

#### Moran's I

Tobler's first law of geography states that "Everything is related to everything else. But near things are more related than distant things." [2]. This observation motivates the examination of spatial autocorrelation. Positive spatial autocorrelation is evident when nearby things tend to be more similar. For example, weather in Los Angeles tends to be more similar to that of San Diego than that of New York. Negative spatial autocorrelation is evident when nearby things tend to be more dissimilar, such as squares on a chessboard. Spatial autocorrelation can arise from an intrinsic process such as diffusion, via communication by physical contact, or as the result of a covariate that has such an intrinsic process. In areal data, data is aggregated over spatial areas such as population in each neighborhood and city. Positive spatial autocorrelation can arise when the areal units of observation are smaller than the scale of the spatial process, such as when the phenomenon occurs at the level of cities while the units of observation are neighborhoods [3].

Moran's I [4] is one of the most commonly used statistics for assessing spatial autocorrelation. It is defined as

$$I = \frac{n}{\sum_{i=1}^{n}\sum_{j=1}^{n} w_{ij}} \cdot \frac{\sum_{i=1}^{n}\sum_{j=1}^{n} w_{ij}\left(x_i-\bar{x}\right)\left(x_j-\bar{x}\right)}{\sum_{i=1}^{n}\left(x_i-\bar{x}\right)^2}, \tag{1}$$

where $w$ is the number of spots or locations, $i$ and $j$ are different locations, or spots in the Visium context, $x$ is a variable with values at each location, and $w_{ij}$ is a spatial weight, which can be inversely proportional to distance between spots or an indicator of whether two spots are neighbors, subject to various definitions of neighborhood. Moran's I is similar to the Pearson correlation between the value at each location and the average value at its neighbors (but not identical, see [5]). Just like Pearson correlation, Moran's I is usually bound between -1 and 1, where positive values indicate positive spatial autocorrelation and negative values indicate negative spatial autocorrelation.

Local Moran's I [6] is defined as

$$I_i = \frac{\left(x_i-\bar{x}\right)\sum_{j=1}^{n} w_{ij}\left(x_j-\bar{x}\right)}{\sum_{i=1}^{n}\left(x_i-\bar{x}\right)^2/(n-1)}, \tag{2}$$

It constitutes an unnormalized and disaggregated form of Moran's I, and describes the contribution of each location to the global Moran's I.

#### MULTISPATI PCA

The Moran's I expression above can be rearranged as

$$I = \frac{1}{\sum_{i=1}^{n}\sum_{j=1}^{n} w_{ij}} \cdot \frac{\sum_{i=1}^{n}\sum_{j=1}^{n} w_{ij}\left(x_i-\bar{x}\right)\left(x_j-\bar{x}\right)}{\sum_{i=1}^{n}\left(x_i-\bar{x}\right)^2/n}, \tag{3}$$

where the denominator is the maximum likelihood estimate (MLE) of the variance of the data. Let $z$ denote the scaled and centered data, whose mean is 0 and variance (using MLE, divide by $n$ instead of $n-1$) is 1. If the spatial weights matrix $W$ is scaled so its rows sum to 1, the denominator of the first term becomes $n$ and Moran's I can be more simply expressed as

$$I = \sum_{i=1}^{n}\sum_{j=1}^{n} w_{ij} z_i z_j / n. \tag{4}$$

This can be written more succinctly as $I = \mathbf{z}^T\mathbf{W}\mathbf{z}/n$. Let $\mathbf{Z}$ denote a matrix whose columns are scaled (divided by the standard deviation, making the variance 1) and centered (subtract the mean, thus summing to 0) variables and whose rows are observations such as cells in space. Then this expression of Moran's I can be generalized to multiple variables: $\mathbf{M} = \mathbf{Z}^T\mathbf{W}\mathbf{Z}/n$. The diagonal of $\mathbf{M}$ is Moran's I coefficients of the variables in $\mathbf{Z}$. We can find the eigenvalues and eigenvectors of this matrix as a spatially informed form of PCA proposed by Wartenberg in 1985 [7].

This is analogous to PCA, in which the covariance matrix $\mathbf{X}^T\mathbf{X}/n$ is diagonalized after each variable (column) in $\mathbf{X}$ is centered. In non-spatial PCA, the first eigenvector (principal component, or PC), which has the largest eigenvalue, finds the linear combination of the original variables that explains the most variance. The corresponding eigenvalue is the amount of variance explained by this PC. The second PC (PC2) is found by maximizing the variance again provided that PC2 is orthogonal to PC1, and so on. Because the covariance matrix is symmetric and positive semi-definite, all of its eigenvalues are real and non-negative, and it has orthogonal eigenvectors, i.e. the eigenvectors are orthogonal to each other, which can be normalized to have norm 1 (i.e. orthonormal). However, the interpretation of the eigenvalues and eigenvector of $\mathbf{M}$ is not explored in reference number [7].

The Wartenberg method summarized above was generalized for the statistical triplet of multivariate data analysis in the duality diagram paradigm in [8], and implemented in the `adespatial` R package as MULTISPATI. With `adespatial`, the spatial information can not only be used for PCA but also for other multivariate analyses with the duality diagram such as correspondence analysis. However, for now, Voyager's much faster implementation of MULTISPATI only applies to PCA. See [9] for an introduction to the duality diagram.

Let $\mathbf{X}$ be a data matrix with $n$ rows and $p$ columns with observations in rows and variables in columns, and assume that each variable is centered. MULTISPATI PCA seeks to find vector $\mathbf{u}_1$ of norm 1 that maximizes $Q(\mathbf{u}_1) = \mathbf{u}_1^T\mathbf{X}^T\mathbf{W}\mathbf{X}\mathbf{u}_1/n$. Let $\mathbf{a}_1 = \mathbf{X}\mathbf{u}_1$. Then MULTISPATI PCA maximizes $Q(\mathbf{u}_1) = \mathbf{a}_1^T\mathbf{W}\mathbf{a}_1/n$. Remember the matrix expression of Moran's I, $I = \mathbf{z}^T\mathbf{W}\mathbf{z}/n$. Because $\mathbf{X}$ is centered, all columns sum to 0, so $\mathbf{1}^T\mathbf{X} = \mathbf{0}^T$, where $\mathbf{1}$ is a vector of $n$ 1's and $\mathbf{0}$ is a vector of $p$ 0's. Hence, $\mathbf{1}^T\mathbf{a}_1 = \mathbf{1}^T\mathbf{X}\mathbf{u}_1 = 0$, meaning that $\mathbf{a}_1$ is also centered. Then we need to scale $\mathbf{a}_1$ so its variance is 1 by dividing it by its standard deviation, which is square root of the variance. The MLE of the variance is $\sum_{i=1}^n a_{1i}^2/n = \|\mathbf{a}_1\|^2/n$, so $I(\mathbf{a}_1) = \mathbf{a}_1^T\mathbf{W}\mathbf{a}_1/\|\mathbf{a}_1\|^2$. Therefore, $Q(\mathbf{u}_1) = I(\mathbf{a}_1)\|\mathbf{a}_1\|^2/n$. Hence for PC1, MULTISPATI PCA maximizes the product of Moran's I of the projection of the observations onto PC1 and variance explained by PC1.

Just as in non-spatial PCA, these maximizations are achieved by diagonalizing the spatially weighted covariance matrix; the eigenvalues are $Q(\mathbf{u}_i)$. Because $\mathbf{W}$ doesn't have to be symmetric, $\mathbf{X}^T\mathbf{W}\mathbf{X}$ doesn't have to be symmetric. So in practice, it's preferable to diagonalize the symmetric matrix $\mathbf{H} = \frac{1}{2n}\mathbf{X}^T(\mathbf{W}^T + \mathbf{W})\mathbf{X}$ instead, which gives the same eigenvalues as $\mathbf{X}^T\mathbf{W}\mathbf{X}/n$; the eigenvalues are guaranteed to be real when computed and the eigenvectors are orthonormal. However, since asymmetric real matrices don't have orthonormal eigenvectors, the eigenvectors of $\mathbf{H}$ are different from those of $\mathbf{X}^T\mathbf{W}\mathbf{X}/n$. The eigenvectors can be interpreted as if a symmetrized spatial weights matrix $(\mathbf{W}^T + \mathbf{W})/2$ is used for the spatially weighted covariance matrix. The effects of different choices of $\mathbf{W}$ on results remains to be investigated.

### Supplementary Note 3: Defaults in compatibility tests

- Because SFE inherits methods from SCE, the R vignettes use `scater` and `scran` [10] from the SCE ecosystem for QC, data normalization, and non-spatial EDA. Data normalization in scater computes $\log_2\left(\frac{x}{N/\bar{N}} + 1\right)$, where N denotes the total UMI count in one Visium spot, $\bar{N}$ is the average total UMI count in all spots in this dataset, and $x$ is the UMI count of one gene in the Visium spot of interest. The log transform has been shown to perform well both for variance stabilization [11] and in preserving the k-nearest-neighbor graph [12]. The pseudocount (default to 1), library size factors (default to $N/\bar{N}$), and transform (default to log2) can be changed. The size factor is centered on 1 to make it easier to translate log normalized counts back to raw counts. Log 2 is used because differences in values can be interpreted as log fold change. The Python implementation uses the same data normalization.

- `scran` finds highly variable genes (HVGs) as follows: with default parameters, a parametric non-linear curve $y = \frac{ax}{x^n+b}$ of variance vs. mean is fit for each gene of the log normalized data. Subsequently, the log ratio of the actual variance to the fitted variance from the curve is calculated, and a LOWESS curve is fitted to this log ratio vs. mean scatter plot for each gene. The "technical" component of the variance is the fitted values from the Lowess curve. The "biological" component is the difference between the actual log ratio and the Lowess fitted log ratio. The top HVGs are genes with the largest biological component. The default parameters are used in R vignettes not covered by compatibility tests. For the basic vignettes covered by compatibility tests, because we did not find a Python implementation of Lowess, the Python version reimplements `scran`'s HVG method when using parameter `lowess = FALSE` in `modelGeneVar()`, i.e. omitting the Lowess step, so the fitted, "technical" values come from the parametric curve and the "biological" component is the difference between the actual variance and the curve. This method assumes that most genes are not biologically interesting to the study of interest.

- Voyager sets the number of top HVGs to 2000 to be consistent with the Seurat convention. This number is arbitrary and may be changed in the future based on benchmarks of various methods [13].

- PCA is performed on log normalized data with the HVGs to be consistent with the Seurat convention. Data is scaled before performing PCA, i.e. each gene from the log normalized data is scaled and centered to have mean 0 and variance 1, so genes that are more abundant but not necessarily more biologically variable don't drown out genes that are less abundant but more biologically variable in the top principal components. This can happen because gene expression data is overdispersed, so the variance not only increases with mean but also exceeds the mean, so the variance is greater than one would expect from a Poisson distribution. The data is scaled also to be consistent with the Seurat convention.

- When scaling the data, the variance is computed, and the data for each gene is divided by the variance. One can either divide by n or (n-1); the former is the default in Numpy, as the maximum likelihood estimate of variance although it's biased, while the latter is the default in R and scanpy, as an unbiased estimate. We divide by (n-1) in both the R and Python Voyager implementations, to be consistent with the R and scanpy convention.
- The number of PCs used for non-spatial clustering is determined by the elbow plot per the Seurat convention. In the vignette using mouse olfactory bulb Visium data, the cell projections into the PCs are also visually inspected to exclude PCs that appear to pick up artifacts and outliers.
- The k in k-nearest-neighbor graph used in Leiden clustering is the default in `igraph`, which is 10.
- Leiden clustering is used because both Leiden and Louvain are conventionally used in scRNA-seq and Leiden has improved upon Louvain.
- In Leiden clustering, the resolution parameter (0.5 in the basic vignettes) and objective function (modularity in the basic vignettes) are chosen to give a few clusters that appear well-separated in the first few PCs but not so many that their colors are difficult to tell apart when plotting. This is for visualization purposes and may not be the best biological choice. However, Voyager does not always produce reproducible Leiden clustering results due to the random nature of the Leiden algorithm. Results can differ with the R and Python implementations despite setting a random seed.
- For the Chromium PBMC dataset, to find the cluster marker genes, `findMarkers()` in `scran` was used, with Wilcoxon rank sum tests, only testing up regulations. The most highly ranked genes are those differentially expressed in the current cluster and all other clusters (`pval.type = "all"`). The top marker gene for each cluster is the one with the smallest p-value; there were no ties in this case. The Voyager Python implementation re-implements the `scran` method to match the results. These parameters were chosen because they are similar to Seurat conventions.
- When applying spatial statistics methods to the k-nearest-graph in the Chromium PBMC dataset, $k = 10$ (not including self) was chosen to be consistent with Leiden, so the spatial statistics can be better compared to Leiden clustering.
- Voyager uses "W" style edge weights for the spatial neighborhood graph, i.e. the rows of the binary adjacency matrix are normalized to sum up to 1. For a k nearest neighbor graph where all nodes have the same degree, all the edge weights are the same, so will not lead to a different Moran's I. However W style was chosen because it's preferable for the Moran scatter plot, which was performed in the vignette. Using W style, the slope of the line fitted to the Moran scatter plot is Moran's I [14]. W style is also the default across Voyager (except that binary style is recommended for Getis-Ord Gi*) because it is the default in spdep and it simplifies the math for some spatial statistics such as Moran's I, Lee's L, and MULTISPATI PCA and simplifies the interpretation of spatial auto-regressive models.
- For local Moran's I, such as shown in Figure 2G, spdep and the PySAL esda package have different defaults. In order for the results to perfectly match, set `mlvar = FALSE` in the R implementation of Voyager, which is passed to `spdep`, so both implementations divide by n-1 when computing the variance.
- The Voyager R and Python implementations use the same palettes. See Supplementary Figure 3 on colorblindness simulations of the default palettes.
- In the Wilcoxon rank sum test comparisons in Supplementary Note 4, for `scran`, non-default `pval.type = "all"` and `direction = "up"` in the `findMarkers()` function were used, because the only DE functionality implemented in VoyagerPy is equivalent to using these parameters in `scran`.

These are defaults specific to the R package and website at present:

- By default, the image, if present, is not plotted behind the geometries of Visium spots or cells, because the geometries cover up much of the image, and the image can distort color perception of the geometries. However, plotting the image can be useful to visually relate a value in space to histology.
- In neighborhood view ESDA, depending on how the spatial graph is defined, sometimes there are cells that don't have any spatial neighbors. The argument `zero.policy` in `spdep` determines what to do with these singleton cells. By default, in `spdep`, the global option is used, and when a global option is absent, `spdep` throws an error when there are singletons. Voyager generally follows the `spdep` default. However, in many examples, `zero.policy` is set to TRUE where singletons can occur, such as in the correlogram when some cells or spots don't have higher order neighbors. When `zero.policy = TRUE`, spatially lagged values of singletons are set to 0, while when it's `FALSE`, the spatially lagged values are `NA`. `TRUE` is chosen to silently drop the singletons when spatially lagged values are needed without stopping the computation for non-singletons or causing the `NA`'s to propagate.
- In vignettes for image based single cell resolution datasets, we used k-nearest-neighbor graph with $k = 5$ for cell centroids, found with the KMKNN algorithm in `BiocNeighbors` (v1.18.0). We chose $k = 5$ because most "well-behaved" real world tessellations of the 2D plane are somewhere between a square ($k = 4$ rook style) and hexagonal ($k = 6$) tessellation [15]; $k = 5$ seems reasonable based on visual inspection. Furthermore, for a larger dataset with over hundreds of thousands of cells, the k nearest neighbor graph is much faster to compute than most other types of neighbors, such as those that require triangulation. Although with GEOS spatial indexing, the polygon contiguity graph is fast to compute for larger datasets, given the imperfection of cell segmentation, many cells that don't appear contiguous in the polygons might in fact be physically touching, resulting into many false negatives in spatial neighbors and many singletons. As a result, we did not use the polygon contiguity graph for the cells. Inverse distance weighting is used since k nearest neighbors may or may not be physically interacting so further cells have less weight. W style edge weight normalization is used, for reasons mentioned above. However, the polygon contiguity graph is used for spatial analyses of attributes of the myofibers in the mouse skeletal muscle Visium dataset, where false negatives are not an issue.

### Supplementary Note 4: Log fold changes in Seurat, scanpy, and scran

#### Introduction

In a recent survey of methods for finding marker genes from single-cell RNA-seq data [16], the authors urge that "extreme care should be taken when comparing the log fold-changes output by Seurat and Scanpy" because the programs are using different formulas for the calculations. Specifically, in the notation of [16], the Seurat calculation is given by

$$R_g = \log_2\left(\frac{1}{n_1}\sum_{i\in G_1}(\exp(Y_{ig}) - 1) + 1\right) - \log_2\left(\frac{1}{n_2}\sum_{i\in G_2}(\exp(Y_{ig}) - 1) + 1\right), \tag{5}$$

where $Y_{ig}$ are the log-transformed expression values for cell $i$ and gene $g$, $G_1$ and $G_2$ are the indices for two groups of cells, and $n_1$ and $n_2$ are the numbers of cells in the respective groups. In contrast, the Scanpy calculation is

$$P_g = \log_2\left(\exp\left(\frac{1}{n_1}\sum_{i\in G_1} Y_{ig}\right) - 1 + \epsilon\right) - \log_2\left(\exp\left(\frac{1}{n_2}\sum_{i\in G_2} Y_{ig}\right) - 1 + \epsilon\right), \tag{6}$$

where $\epsilon = 10^{-9}$. The different results produced by these formulas (see Supplementary Figure 8 for an example) raises the question of which is the correct one. We examine this question, and find that errors in arithmetic, problems with statistical reasoning, absence of biophysics considerations, and lack of biological motivation all contribute to discrepancies in the formulas and confusion for users.

Before delving into the formulas, we review a handful of elementary properties of the logarithm. We include a review of them because in our experience, users of Seurat and Scanpy are not always familiar with these properties due to a lack of training in mathematics for biology students [17]. However, an understanding of the properties of logarithm is essential for understanding why one log-fold change formula may be preferred over another.

The logarithm $\log(x)$ is defined for real numbers $x > 0$. Unless otherwise specified, $\log(x)$ refers to the natural log of $x$ (sometimes also denoted $\ln(x)$). Thus, if $y = \log(x)$ then $x = \mathrm{e}^y$, sometimes also denoted $y = \exp(x)$. The notation $\log_2(x)$ means that the logarithm is computed with respect to base 2, i.e. if $y = log_2(x)$ then $x = 2^y$. A *pseudocount* is the number $\epsilon$ in the expression $\log(x + \epsilon)$. The logarithm of a product $\log(xy)$ is the sum of the logarithms $\log(x)$ and $\log(y)$, i.e. $\log(xy) = \log(x) + \log(y)$. Also, $\log(x^a) = a\log(x)$, and in particular, $\log(\frac{1}{x}) = -\log(x)$. These properties imply that $\log(\frac{x}{y}) = \log(x) - \log(y)$, which is central to the computation of log-fold change. The power property of logarithms implies that the logarithm of the geometric mean of numbers $x_1, \ldots, x_n$ is $\log((x_1 \cdots x_n)^{\frac{1}{n}}) = \frac{1}{n}\sum_{i=1}^{n}\log(x_i)$. The arithmetic mean of $x_1, \ldots, x_n$ is defined to be $\frac{1}{n}\sum_{i=1}^{n} x_i$, and therefore the logarithm of the geometric mean is the arithmetic mean of $\log(x_1), \ldots, \log(x_n)$. The arithmetic mean and geometric mean are related via an inequality: $(x_1 \cdots x_n)^{\frac{1}{n}} \leq \frac{1}{n}\sum_{i=1}^{n} x_i$ with equality attained if, and only if, $x_1 = x_2 = \cdots = x_n$. The logarithm is frequently used in the analysis of count data, because it provides a good approximation to the inverse hyperbolic sine function, and is therefore near optimal for variance stabilization of negative binomially distributed data [18].

#### Scaling

Let $c(i)$ be the cluster to which cell $i$ belongs. The model then is that the count $X_{ig}$ for gene $g$ in cell $i$ is drawn from a (negative binomial?) distribution with mean $s_i\mu_{c(i)g}$ where $s_i$ is a scaling factor for cell $i$. The desired log-fold-change between clusters $j$ and $k$ is then $\log(\mu_{jg}) - \log(\mu_{kg})$. If $s_i$ were uniformly 1 then $\frac{1}{n_j}\sum_{i;c(i)=j} X_{ig}$ would be an estimator of $\mu_{jg}$ but it's not entirely clear that using $\frac{1}{n_j}\sum_{i;c(i)=j} X_{ig}/s_i$ as an estimator is really valid in the general case.

#### Seurat

We begin by noting that the value $Y_{ig}$ that appears in the Seurat formula $R_g$ is not the raw count of gene $g$ in cell $i$. Rather $Y_{ig}$ is typically obtained by dividing the raw count $X_{ig}$ by the total number of counts in cell $i$, and then multiplying by the number $10,000$ (users may change this by adjusting options). Specifically, $Y_{ig} = \log(\frac{X_{ig}}{S_i} + 1)$ where $S_i = \frac{\sum_g X_{ig}}{10{,}000}$. Thus, $\exp(Y_{ig}) - 1 = \frac{X_{ig}}{S_i}$, and $\frac{1}{n_1}\sum_{i \in G_1}(\exp(Y_{ig}) - 1)$ (respectively $\frac{1}{n_2}\sum_{i \in G_2}(\exp(Y_{ig}) - 1)$) is the arithmetic mean of $\frac{X_{ig}}{S_i}$ computed across the cells in $G_1$ (respectively $G_2$).

The sample mean $\frac{1}{n_1}\sum_{i \in G_1} X_{ig}$ (in what follows we focus on $G_1$, although the claims and results hold for computations with respect to $G_2$ as well), makes sense to compute, as it can be understood to be the maximum likelihood estimate (MLE) for the mean of a negative binomial distribution, namely the distribution for the molecule counts $X_{ig}$. The modeling of the $X_{ig}$ with a negative binomial distribution represents an assumption about the data in an experiment, but is justifiable [19]. The arithmetic means computed in $R_g$ are not, however of the $X_{ig}$. As noted above, they are of $\frac{X_{ig}}{S_i}$, whose distribution will depend on the cell depths $S_i$; moreover the values $\frac{X_{ig}}{S_i}$ are not integers. Thus, the use of the MLE for the negative binomial distribution may not be appropriate, although in practice the log-fold change is computed for cells in two distinct cell types, and therefore the sets $G_1$ and $G_2$ will be homogeneous leading to $\frac{X_{ig}}{S_i}$ being (approximately) a constant multiple of the $X_{ig}$. When this is the case, the sample mean $\frac{1}{n_1}\sum_{i \in G_1} X_{ig}$ is likely to yield a good estimate of the scaled mean for the negative binomial distribution of the $X_{ig}$.

The next piece in the Seurat formula $R_g$ is the use of a pseudocount (+1) when computing the logarithm of the mean estimate. This pseudocount is presumably present to avoid an error from trying to compute the logarithm of zero when a gene has no counts in one of the groups $G_1$ or $G_2$. The Seurat authors seem to have borrowed this "fix" from the Seurat logarithm normalization procedure, which also uses a pseudocount of +1 in the computation of the values $Y_{ig}$, i.e., $Y_{ig} = \log(\frac{X_{ig}}{S_i} + 1)$. In the case of normalization of negative binomially distributed data, the use of a pseudocount amounts to an assumption about the extent of overdispersion in $X_{ig}$ with respect to the Poisson distribution [20]. Specifically, the logarithm makes sense

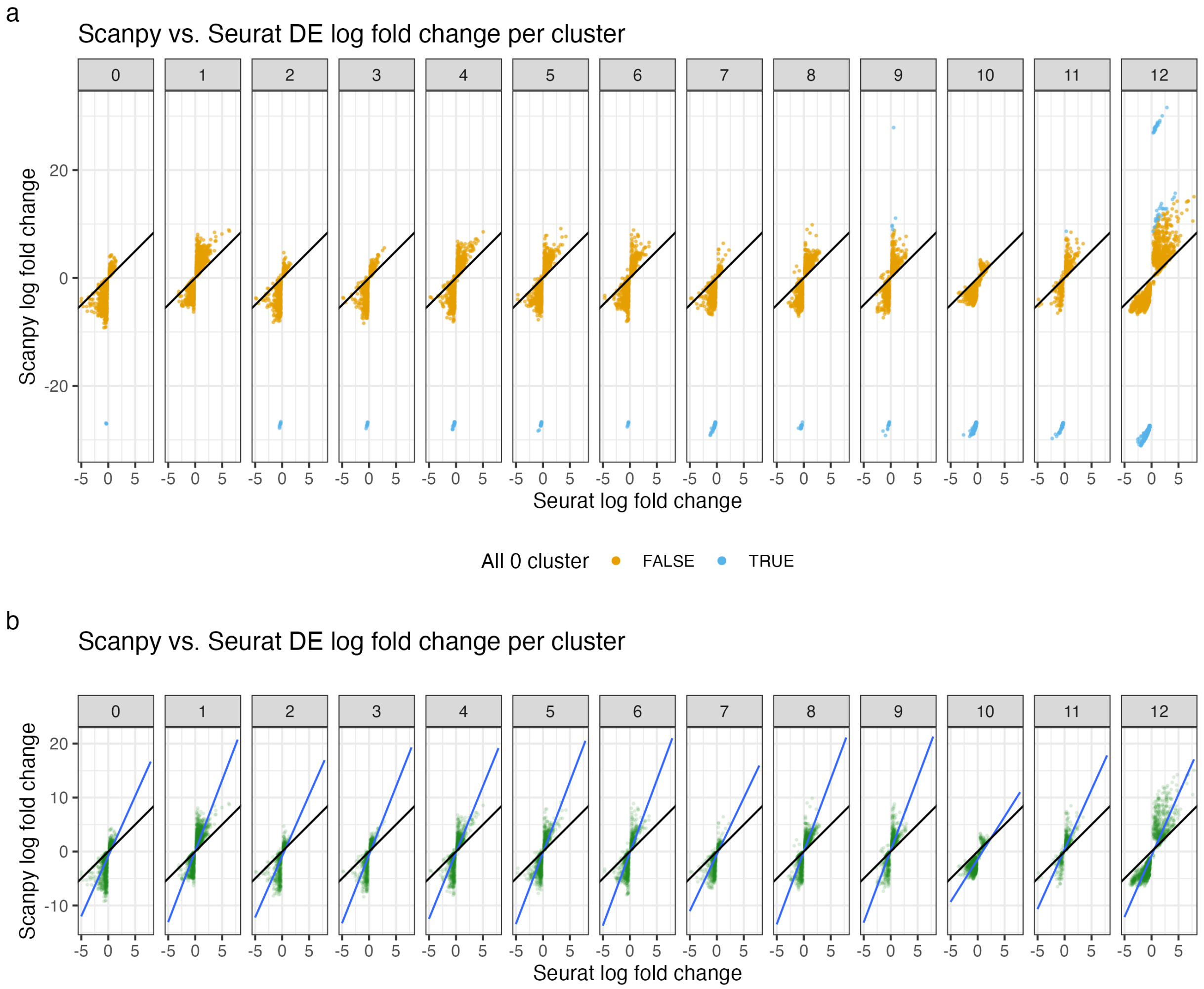


Supplementary Figure 8: Comparisons between Seurat and Scanpy log fold changes in each cluster of a PBMC 5k dataset. a) Scanpy log fold change of each gene in each cluster vs. the rest of the cells is plotted on the y-axis, and the x-axis is the Seurat log fold changes (see Methods). Each point is a gene, and genes that are not detected (all 0) either in the cluster of interest or in all cells outside the cluster (complement) are colored blue. b) Same as panel a, but without genes that are not detected in any cluster or their complements. The black lines are $y = x$, and the blue lines are least square fitted independently for each cluster and have a slope greater than 1 due to the large pseudocount in Seurat discussed in the text.

as a variance stabilizing transformation for negative binomially distributed data, and as discussed above, the negative binomial assumption makes sense for the $X_{ig}$. A pseudocount of $+1$ corresponds to an overdispersion estimate of the form $\sigma^2 = \mu + 0.5\mu^2$ where $\sigma^2$ is the variance of the negative binomial distribution and $\mu$ the mean [20, 21]. However, none of this is relevant when computing the log-fold change, where the use of the logarithm is unrelated to variance stabilization, and instead is just a way to symmetrize fold-change around zero. Unfortunately, the use of a large pseudocount in the computation of $R_g$ reduces and distorts the log-fold change (Supplementary Figure 8), especially for low-expressed genes. This partly explains why, in Supplementary Figure 8, for almost all genes, the Scanpy estimated log-fold change is higher than the Seurat log-fold change.

#### Scanpy

The Scanpy formula $P_g$ is also based on on expression estimates $Y_{ig}$ that are computed according to user specifications; similarly to Seurat, the Scanpy primary tutorial demonstrates the computation of the $Y_{ig}$ as $Y_{ig} = \log(\frac{X_{ig}}{S_i} + 1)$. Thus, $\frac{1}{n_1}\sum_{i \in G_1} Y_{ig}$ is the logarithm of the geometric mean of $\exp(Y_{ig})$, i.e $\log((\prod_{i \in G_1}(\frac{X_{ig}}{S_i}+1))^{\frac{1}{n}})$. The log of geometric mean is the MLE for the mean of log-normal distributed data, so it makes sense to use the geometric mean if one assumes that the $\exp(Y_{ig})$ are log-normally distributed, however the authors of Scanpy explain that this is not the case, writing in [22] that "scRNA-seq data are not in fact log-normally distributed". Moreover, there is an arithmetic error in the computation of $P_g$, evident in the subtraction of 1 after computing the geometric mean in the $P_g$ formula $\log_2\left(\exp\left(\frac{1}{n_1}\sum_{i \in G_1} Y_{ig}\right) - 1 + \epsilon\right)$. The subtraction is intended to adjust for the fact that the geometric mean is computed for $\frac{X_{ig}}{S_i} + 1$ and not $\frac{X_{ig}}{S_i}$; presumably this was a matter of convenience as the $Y_{ig}$ are stored in the `anndata` object that Scanpy uses for other purposes. However, while the arithmetic mean is linear, the geometric mean is not, and in general $((x_1 + 1)\cdots(x_n + 1))^{\frac{1}{n}} - 1 \neq (x_1 \cdots x_n)^{\frac{1}{n}}$. Again, in most cases, the homogeneity of the groups $G_1$ and $G_2$ comes to the rescue as $((x_1 + 1)\cdots(x_n + 1))^{\frac{1}{n}} - 1 \approx (x_1 \cdots x_n)^{\frac{1}{n}}$ when the $x_i$ are close to each other. The formula $P_g$ also includes a pseudocount to avoid an error in the case when the $Y_{ig}$ are all equal to zero for a group (and thus the attempted evaluation of the logarithm of zero), but in the case of $P_g$ the pseudocount used is $\epsilon = 10^{-9}$, thus avoiding the problems with the +1 pseudocount in $R_g$.

The $\epsilon = 10^{-9}$ leads to the outlying blue clusters around ±30 on the y-axis in Supplementary Figure 8A, because when a gene is all 0 in a cluster or its complement, one of the two terms in Equation 2 becomes $\pm\log_2 10^{-9} \approx \pm 29.90$. When the cluster itself ($G_1$) is all 0 but not its complement ($G_2$), the log-fold change is -30; when $G_2$ is all 0 but not $G_1$, the log-fold change is around 30. However, by the definition of log-fold change $\log(\mu_{jg}) - \log(\mu_{kg})$, the log-fold change should really be $-\infty$ and $\infty$ respectively. In contrast, Seurat does not give a large value of log-fold change in the all 0 cases. When the gene is all 0 in all cells in $G_2$, the second term of Equation 1 becomes $\log_2 1 = 0$, so the log-fold change becomes $\log_2\left(\frac{1}{n_1}\sum_{i \in G_1}(\exp(Y_{ig}) - 1) + 1\right)$.

In addition, note that in the standard Scanpy workflow such as indicated in its tutorials, prior to principal component analysis (PCA), the gene count matrix is scaled after log normalization so each gene has mean 0 and variance 1. The scaling was not performed in [16]. When the data is scaled, the Wilcoxon rank sum test is unaffected as it is based on ranks. However, because scaling introduces negative values, the log-fold change cannot be computed. As the DE output of both Seurat and Scanpy contains p-values and users may use the adjusted $p < 0.05$ cutoff to select genes for further analyses, we compare the p-values to compare the DE results from the standard workflow (Supplementary Figure 9A). Despite using the more conservative Bonferroni correction, Seurat tends reports more significant p-values than Scanpy, which uses the Benjamini-Hochberg correction. In contrast, p-values reported by VoyagerPy and `scran` largely agree, except for some genes with less significant p-values for clusters 11 and 12.

#### Scran

Unlike Seurat and Scanpy, `scran` does not compute the log-fold change as the effect size when a Wilcoxon rank sum test is performed for differential expression (DE). Rather, the area under the receiver operating characteristic curve (AUC) is used as the effect size. While the `scran` package computes log-fold change for other DE tests, VoyagerPy only implements the Wilcoxon rank sum test for DE according to the implementation in `scran`. Hence the compatibility test only concerns the AUC. In contrast to the log-fold changes in Seurat vs. Scanpy, the AUC's from VoyagerPy and `scran` largely match though there are some discrepancies in genes with lower AUC's (Supplementary Figure 10).

However, `scran` computes the log-fold change for the t-test and binomial test, and the implementation is different for the two tests. By default, the log normalized data is used in DE and is used to compute log-fold changes. Log normalization in `scran` is different from that in Seurat and Scanpy: let $N_i = \sum_g X_{ig}$ denote the total UMI count in cell $i$, and $\bar{N}$ the average total count across all cells in the dataset. By default, the size factor $S_i = N_i/\bar{N}$, so $S_i$ is centered at 1, making it relatively easy to relate $Y_{ig}$ back to $X_{ig}$ because $X_{ig} \approx 2^{Y_{ig}} - 1$. The log normalized count $Y_{ig} = \log_2\left(\frac{X_{ig}}{S_i} + 1\right)$. For the t-test, the log-fold change is

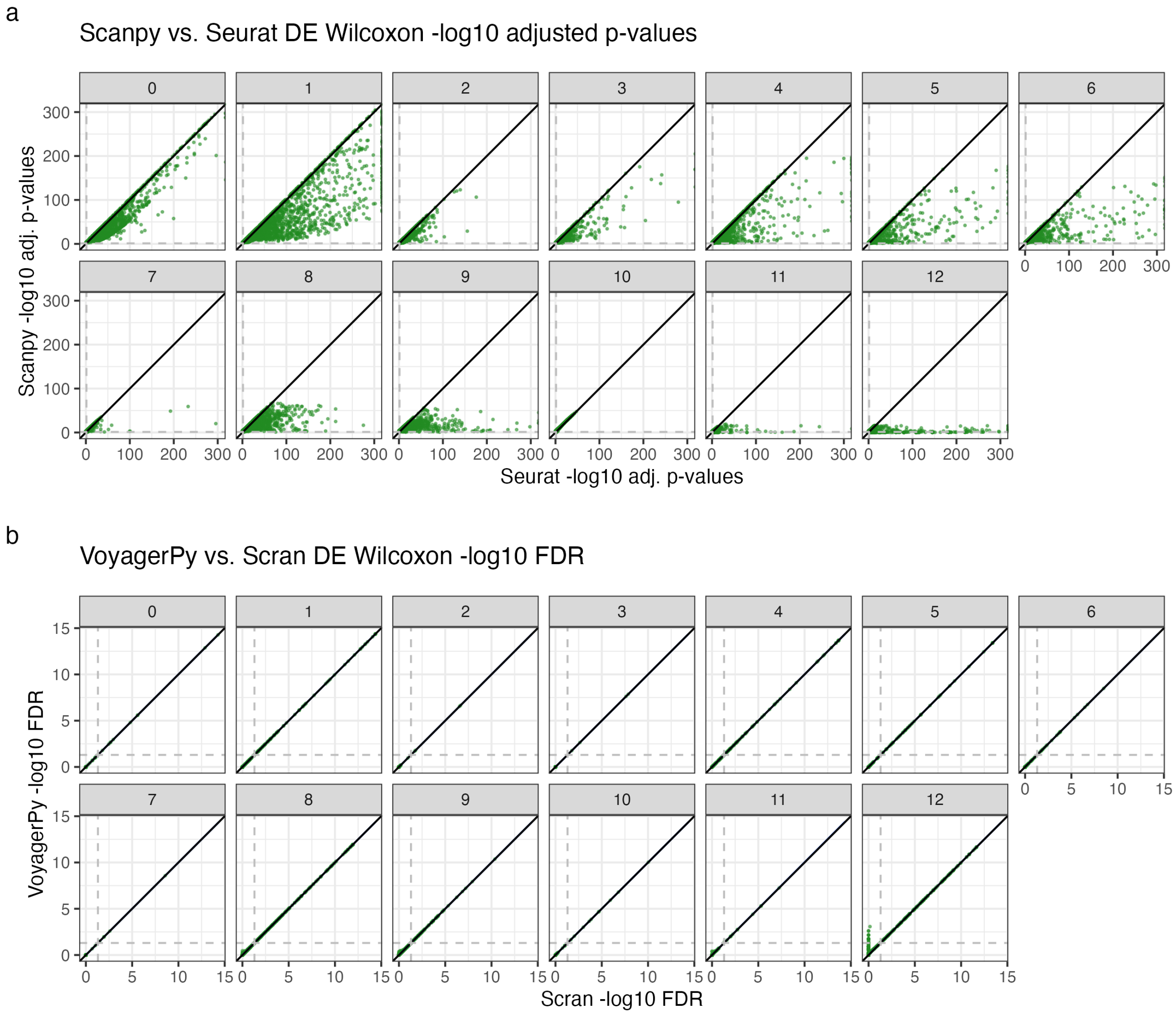


Supplementary Figure 9: Comparisons of adjusted p-values from different implementations of the Wilcoxon rank sum test. a) Comparison between adjusted p-values from Seurat and Scanpy for each cluster; each point is a gene, and the black line is $y = x$, indicating agreement between the two implementations. Dashed horizontal and vertical lines indicate adjusted $p < 0.05$. Seurat reports $p = 0$ for some genes, which are not plotted because $-log_{10}p = \infty$. b) Same as panel a but for VoyagerPy and `scran`.

$$T_g = \frac{1}{n_1}\sum_{i\in G_1} Y_{ig} - \frac{1}{n_2}\sum_{i\in G_2} Y_{ig} = \log_2\left[\left(\prod_{i\in G_1}\left(\frac{X_{ig}}{S_i}+1\right)\right)^{\frac{1}{n_1}}\right] - \log_2\left[\left(\prod_{i\in G_2}\left(\frac{X_{ig}}{S_i}+1\right)\right)^{\frac{1}{n_2}}\right], \quad (7)$$

which is effectively the log-fold change in the geometric means of scaled raw counts with pseudocount in the two clusters. This has the same problems as the Scanpy implementation.

Log-fold change for the binomial test is very different. According to `scran` documentation for `pairwiseBinom`:

> Effect sizes for each comparison are reported as log2-fold changes in the proportion of expressing cells in one group over the proportion in another group. We add a pseudo-count that squeezes

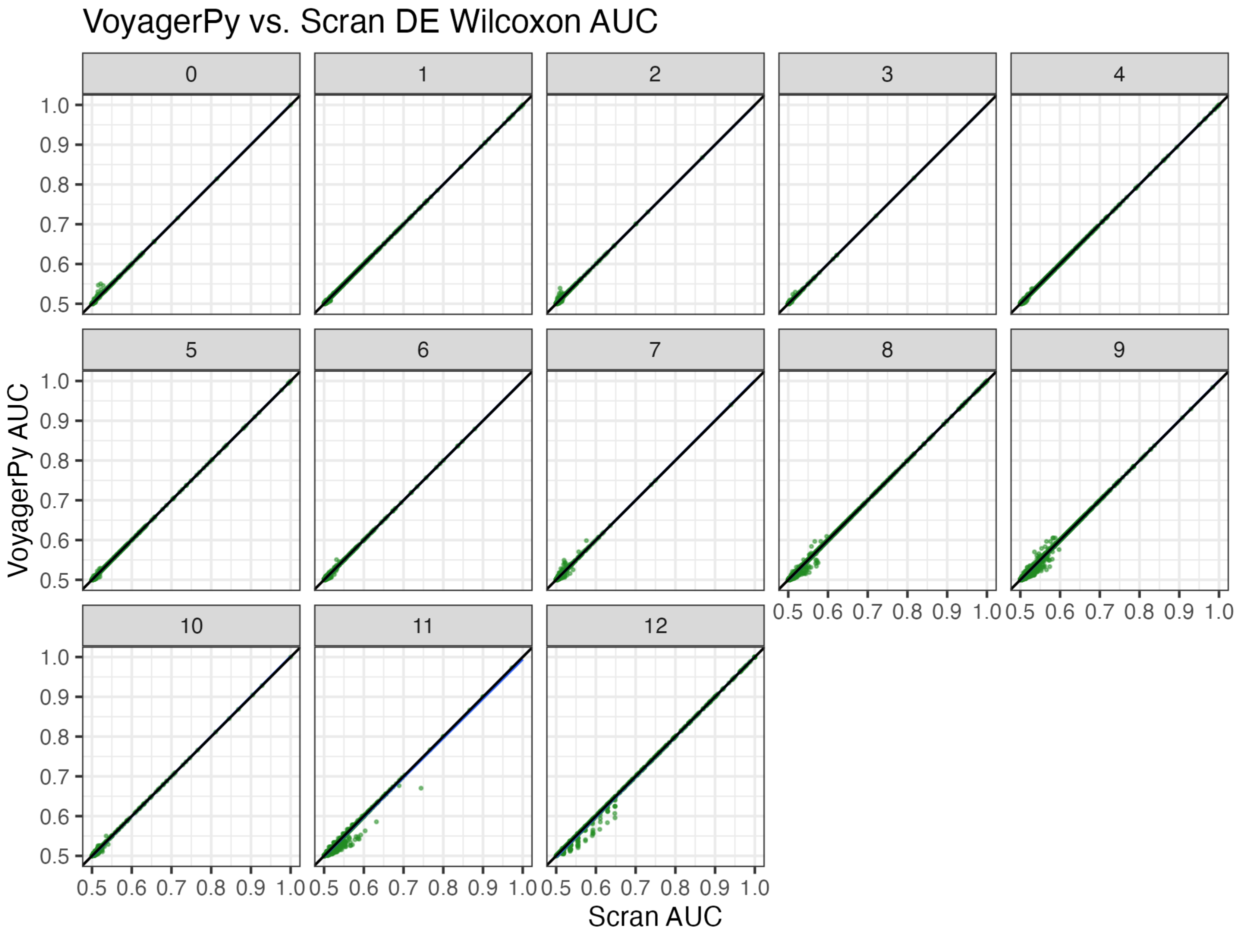


Supplementary Figure 10: Comparison between AUC from VoyagerPy and `scran` from the Wilcoxon rank sum test. Only genes with AUC > 0.5 (i.e. baseline of random guess) are shown. The black line is $y = x$.

the log-FCs towards zero to avoid undefined values when one proportion is zero. This is closely related to but somewhat more interpretable than the log-odds ratio, which would otherwise be the more natural statistic for a proportion-based test.

#### Discussion

There are some assumptions implicit in the discussions above that deserve further scrutiny. First, the assumption that the expression estimates $Y_{ig}$ derived from the counts $X_{ig}$ represent accurate measures of expression is frequently taken for granted, but is not self evident. In practice there is no consensus on how the counts $X_{ig}$ should be obtained [23, 24]. In particular, the pre-processing of single-cell RNA-seq data requires making choices about whether to include in $X_{ig}$ counts of molecules that are from nascent transcripts, or of molecules that are ambiguous as to their origin from mature or nascent transcripts. These issues are particularly vexing when working with single-nuclear RNA-seq [25]. One approach to "integrating" counts of nascent and mature molecules is to use them together to parameterize models of transcription [26], raising the question of whether comparisons of parameter estimates in such models are more suitable for assessing differences in transcription between cell types rather than log-fold change estimates based on the $Y_{ig}$. Moreover, even if one accepts that the (scaled) $Y_{ig}$ are relevant for measuring expression differences between cell types, it may be that the instability of log-fold change with low expression estimates, which are the norm in single-cell RNA-seq experiments due to the sparsity of data, make them inappropriate as proxies for effect sizes [27]. Finally, implicit in all of the above is the emphasis on effect size, frequently in addition to but sometimes in lieu of the use of $p$-values to assess statistical significance. The importance of effect size thresholding in biology research may reflect skepticism of $p$-values, but may also be popular due to the lack of standards for thresholding allowing for the tuning of thresholds to achieve desired results. Sometimes different thresholds are used within a single paper [28].

In summary, there are several problems with the current implementations of the log-fold change calculation in widely used single-cell RNA-seq software. The differences resulting from these issues are non-trivial, and likely affect results reported in the many papers that depend on Scanpy or Seurat. In the Voyager R workflow, the `scran` package is used for many non-spatial analyses. In reimplementing certain `scran` functionalities in VoyagerPy, we have attempted to mitigate this discrepancy for users of Voyager.

#### Data and code availability

The Seurat log-fold change method is described in the source code here.
The Scanpy log-fold change method is described in the source code here.
The `scran` AUC method is implemented here.
The `scran` log-fold change method for t-test is described in the source code here.
The `scran` log-fold change method for binomial test is described in the source code here.
