## Supplementary tables and figures for "Voyager: exploratory single-cell genomics data analysis with geospatial statistics"

#### Supplementary Table 1: Functionalities of Voyager and similar EDA frameworks

Color code: Univariate global, univariate local, categorical, bivariate global, bivariate local, multivariate

| Tradition | Method | Voyager | squidpy | Seurat | Giotto | semla |
| --- | --- | --- | --- | --- | --- | --- |
| Areal, neighborhood view | Moran's I | ✅ | ✅ | ✅ | ✅ | ✅[1] |
| | Geary's C | ✅ | ✅ | | ✅ | |
| | Mantel-Hubert spatial general cross product statistic | ✅ | | | | |
| | Global G | ✅ | | | | |
| | Moran's I, Geary's C analytical and permutation tests | ✅ | ✅ | | ✅ | |
| Geostatistical, distance view | Variogram | ✅ | | | | |
| | Anisotropic variogram | ✅ | | | | |
| | Variogram map | ✅ | | | | |
| Other | Spatially variable genes from binarized data (binSpect) | | | | ✅ | |
| | Wrapper of spatially variable gene methods, e.g. SpatialDE, trendsceek, SPARK, sepal | | ✅ | | ✅ | |
| Areal, neighborhood view | Local Moran's I | ✅ | | | ✅ | |
| | Local Geary's C | ✅ | | | | |
| | Getis-Ord Gi(*) | ✅ | | | ✅ | ✅ |
| | Local spatial | ✅ | | | | |

[1] Semla uses Pearson correlation between the value and spatially lagged value for spatial autocorrelation, which is similar but not exactly the same as Moran's I. See [1].

| Tradition | Method | Voyager | squidpy | Seurat | Giotto | semla |
|---|---|---|---|---|---|---|
| | heteroscedasticity (LOSH) | | | | | |
| | Moran scatter plot | ✅ | | | | |
| | Local Moran's I, Geary C, Getis-Ord Gi(*), LOSH permutation test | ✅ | | | | |
| | LOSH chi-square test | ✅ | | | | |
| Spatial point process | Cross type Ripley's L/F/G function | | ✅ | | | |
| | Spatially variable genes from mark variogram | | | ✅ | | |
| Neighborhood view | Newman's Assortativity | 🟡[2] | | | | ✅ |
| | Neighborhood cell type enrichment (related to join count statistic) | | ✅ | | ✅ | ✅ |
| | Hidden Markov Random Field spatial clustering | | | | ✅ | |
| Other | DE from cell type interaction | | | | ✅ | |
| | Cluster co-occurance score at different distance thresholds | | ✅ | | | |
| | Non-negative least squared cell type deconvolution | | | | | ✅ |
| | DWLS cell type | | | | ✅ | |

[2] Similar analyses have been performed in the Voyager workflow in Supplementary Figures 5-6 with Concordex, although this is not implemented in Voyager itself. Voyager is not meant to be a comprehensive and exclusive spatial data analysis framework; rather it aims to bring the ESDA tradition to complement existing tools, made easier by using the SCE and AnnData classes.

| Tradition | Method | Voyager | squidpy | Seurat | Giotto | semla |
|---|---|---|---|---|---|---|
| | deconvolution | | | | | |
| | Cell-Cell communication scores from ligand-receptor pairs | | ✅ | | ✅ | |
| Areal, neighborhood view | Lee's L | ✅ | | | ✅ | |
| | Lee's L analytical and permutation tests | ✅ | | | | |
| Geostatistical, distance view | Cross variogram | ✅ | | | | |
| | Cross variogram map | ✅ | | | | |
| Areal, neighborhood view | Local Lee's L | ✅ | | | | |
| | Local bivariate Moran's I | ✅ | | | | |
| Areal, neighborhood view | MULTISPATI PCA | ✅ | | | | |
| | Multivariate local Geary's C (permutation test) | ✅ | | | | |

#### Supplementary Table 2: Geometries in SFE and data structures underlying similar EDA frameworks

| Functionality | SFE | squidpy | Seurat | Giotto | Staffli[3] |
|---|---|---|---|---|---|
| Cell/spot centroid coordinates | ✅ | ✅ | ✅ | ✅ | ✅ |
| colGeometries | ✅ | 🟡[4] | ✅[5] | ✅ | |
| rowGeometries | ✅ | | | ✅ | |

[3] Not equivalent to SpatialExperiment. Staffli is a class to hold Visium image and coordinate information, used in conjunction with Seurat objects in the semla package.
[4] Squidpy can perform cell segmentation, but the results are stored as an image layer.
[5] Only works for polygons, which suffices in most cases, while any type of geometry can in theory be used in SFE.

| | | | | | |
|---|---|---|---|---|---|
| annotGeometries | ✅ | | | | |
| Images | ✅ | ✅ | ✅ | ✅ | ✅ |
| Geometric operations | Everything supported by sf and GeoPandas:<br><br>Geometric predicates: Intersect, disjoint, covered by, touch, etc.<br><br>Geometric operations: find intersection,find excluded regions, union, find area, find distance, buffer, find bounding box, spatial joins, simplify geometries, etc.<br><br>Extraction of raster values with vector geometries in the terra R package. | | | | |

#### Supplementary Table 3: Methods to find spatial neighborhood graphs in SFE and similar EDA frameworks

To find spatial graphs in the histological space for neighborhood view spatial analyses, rather than gene expression or PCA space as in graph-based clustering.

| Method | SFE | squidpy | Seurat | Giotto | STUtility/ semla |
|---|---|---|---|---|---|
| Visium spot adjacency | ✅ | ✅ | | | |
| Delaunay triangulation | ✅ | ✅ | | ✅ | |
| K nearest neighbors | ✅ | ✅ | | ✅ | ✅ |
| Distance based neighbors | ✅ | ✅ | | | |
| Gabriel (prune | ✅ | | | | |

| | |
|---|---|
| triangulation) [2] | |
| Relative neighbors (prune triangulation) [3] | ✅ |
| Sphere of influence (prune triangulation) [4] | ✅ |
| Polygon contiguity | ✅ |

### Supplementary Figures

Supplementary Figures 8-10 are part of the Supplementary Notes. The other supplementary figures are in this file.

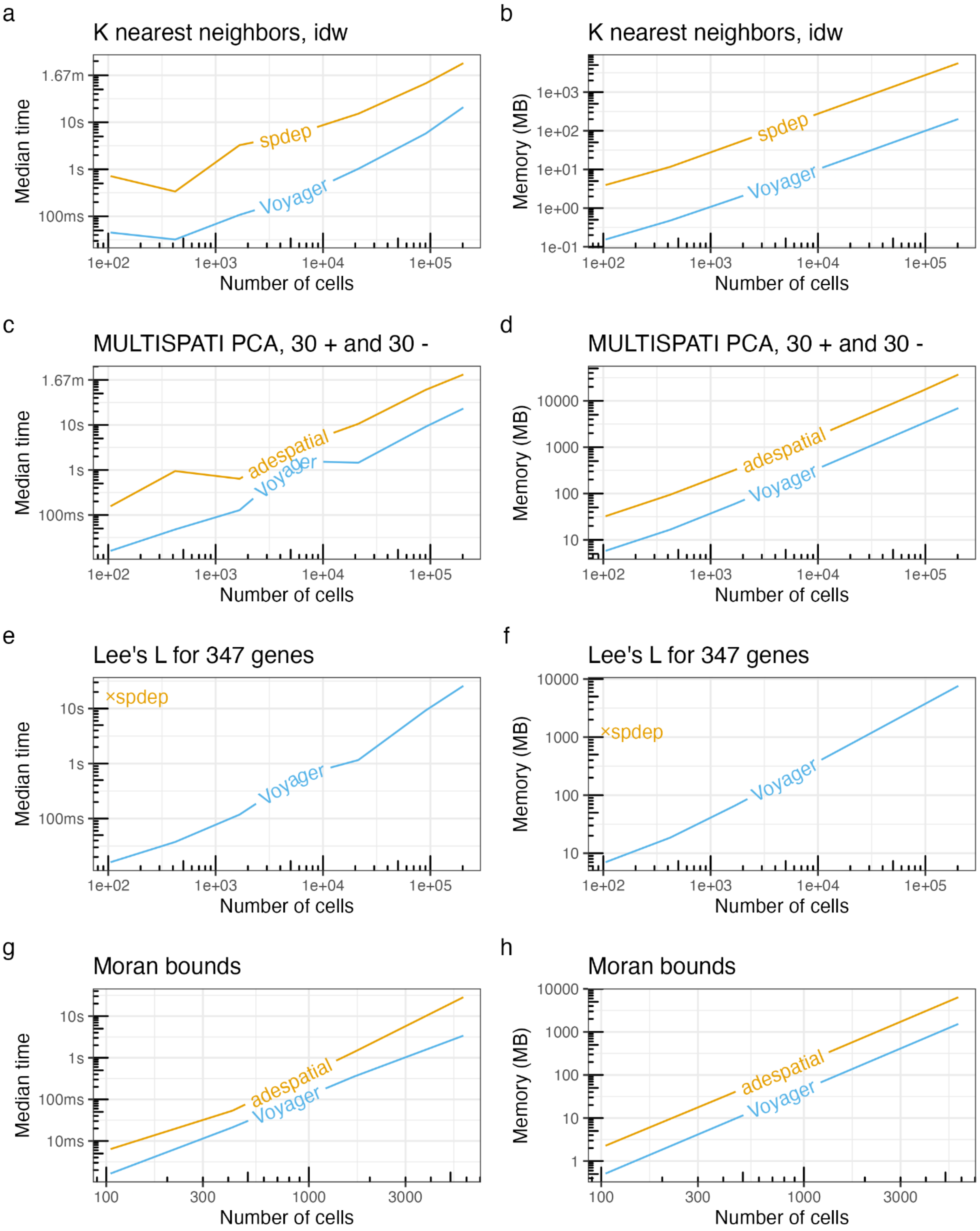

Supplementary Figure 1: Benchmarks of time and memory use of the original implementations and the more efficient implementations in Voyager. A-B) K nearest neighbor graph with inverse distance weighting (idw), with k = 5, W style. C-D) MULTISPATI PCA with 30 positive and 30

negative eigenvalues, using the same k nearest graphs from A-B. E-F) Lee's L for 347 genes. G-H) Finding bounds of Moran's I given spatial neighborhood graph (same as in A-B).

a knn with idw, spdep

| Code | File | Memory (MB) | | Time (ms) |
|---|---|---|---|---|
| ▼ profvis::profvis | | 0 | 63.6 | 1680 |
| ▼ findSpatialNeighbors | AllGenerics.R | 0 | 60.0 | 1580 |
| ▼ .local | | 0 | 60.0 | 1580 |
| ▼ .comp_graph_sample | graph_wrappers.R | 0 | 60.0 | 1580 |
| ▼ .nb2listwdist2.default | graph_wrappers.R | 0 | 54.6 | 1320 |
| ▼ nb2listwdist | graph_wrappers.R | 0 | 54.6 | 1320 |
| ▶ [.sfc | | 0 | 30.4 | 730 |
| ▶ st_distance | | 0 | 22.0 | 540 |
| [ | | 0 | 1.0 | 30 |
| mode<- | | 0 | 0.6 | 10 |
| ▶ .get_centroids | graph_wrappers.R | 0 | 2.5 | 140 |
| ▼ <Anonymous> | | 0 | 2.1 | 80 |
| ▶ knn2nb | graph_wrappers.R | 0 | 1.3 | 60 |
| ▼ knearneigh | graph_wrappers.R | 0 | 0.8 | 20 |
| dbscan::kNN | | 0 | 0.8 | 20 |

b knn with idw, BiocNeighbors

| Code | File | Memory (MB) | | Time (ms) |
|---|---|---|---|---|
| ▼ profvis::profvis | | 0 | 2.4 | 90 |
| ▼ findSpatialNeighbors | AllGenerics.R | 0 | 1.8 | 80 |
| ▼ .local | | 0 | 1.8 | 80 |
| ▼ .comp_graph_sample | graph_wrappers.R | 0 | 1.8 | 80 |
| ▼ <Anonymous> | | 0 | 1.8 | 70 |
| ▼ .knn_bioc | graph_wrappers.R | 0 | 1.8 | 70 |
| ▶ lapply | graph_wrappers.R | 0 | 0.6 | 40 |
| ▶ findKNN | | 0 | 0.3 | 10 |
| ▶ asplit | graph_wrappers.R | 0 | 0.5 | 10 |
| nn$distance[i,] <- nn$distance[i,][ord[[i]]] | graph_wrappers.R | 0 | 0.4 | 10 |
| ▶ colnames | graph_wrappers.R | 0 | 0 | 10 |
| .Call | | 0 | 0.6 | 10 |

c MULSITPATI PCA, adespatial

| Code | File | Memory (MB) | | Time (ms) |
|---|---|---|---|---|
| ▼ profvis::profvis | | 0 | 1013.5 | 1840 |
| ▼ calc_multispati_ade | | 0 | 871.2 | 1260 |
| ▼ dudi.pca | | 0 | 563.5 | 770 |
| ▶ sweep | | 0 | 271.0 | 380 |
| ▼ as.dudi | | 0 | 117.5 | 210 |
| ▶ sweep | | 0 | 82.3 | 110 |
| crossprod | | 0 | 0.9 | 40 |
| eigen | | 0 | 3.3 | 30 |
| as.matrix.data.frame | | 0 | 15.6 | 10 |
| ▶ apply | | 0 | 161.8 | 170 |
| ▼ multispati | | 0 | 231.2 | 390 |
| ▶ Ops.data.frame | | 0 | 82.2 | 150 |
| ▶ apply | | 0 | 84.1 | 140 |
| t.default | | 0 | 18.1 | 30 |
| as.matrix.data.frame | | 0 | 33.2 | 30 |
| eigen | | 0 | 4.2 | 20 |
| ▶ data.frame | | 0 | 9.4 | 20 |
| as.data.frame.matrix | | 0 | 61.1 | 60 |
| ▶ as.matrix | | 0 | 0 | 20 |
| t.default | | 0 | 15.3 | 20 |

d MULSITPATI PCA, Voyager

| Code | File | Memory (MB) | | Time (ms) |
|---|---|---|---|---|
| ▼ profvis::profvis | | 0 | 162.3 | 230 |
| ▼ multispati_rsp | | 0 | 158.6 | 200 |
| ▶ sweep | SFEMethod-multivariate.R | 0 | 61.4 | 90 |
| %*% | | 0 | 47.3 | 80 |
| ▶ colVars | | 0 | 30.7 | 10 |
| t.default | | 0 | 16.6 | 10 |
| ▶ + | | 0 | 2.6 | 10 |
| <Anonymous> | | 0 | 0.9 | 20 |
| %*% | SFEMethod-multivariate.R | 0 | 2.8 | 10 |

e Lee's L, spdep, 105 cells

| Code | File | Memory (MB) | | Time (ms) |
|---|---|---|---|---|
| ▼ profvis::profvis | | -1293.0 | 1274.4 | 21300 |
| ▼ run_pairwise_L | | -1244.2 | 788.4 | 14740 |
| ▼ spdep::lee | | -1244.2 | 729.8 | 13660 |
| ▼ lag.listw | | -369.0 | 263.3 | 5360 |
| stopifnot | | -369.0 | 44.5 | 1940 |
| get.listw_is_CsparseMatrix_Option | | 0 | 30.4 | 460 |
| card | | 0 | 23.9 | 350 |
| $ | | 0 | 17.1 | 180 |
| length | | 0 | 9.2 | 80 |
| lapply | | -427.1 | 232.6 | 3550 |
| mean | | 0 | 32.9 | 490 |
| mean.default | | 0 | 21.6 | 330 |
| stopifnot | | 0 | 15.1 | 260 |
| unlist | | 0 | 16.7 | 240 |
| <GC> | | -386.0 | 0.7 | 200 |
| $ | | 0 | 9.9 | 90 |
| length | | 0 | 7.8 | 80 |

f Lee's L, Voyager, 5700+ cells

| Code | File | Memory (MB) | | Time (ms) |
|---|---|---|---|---|
| ▼ profvis::profvis | | 0 | 218.3 | 280 |
| ▼ calculateBivariate | AllGenerics.R | 0 | 156.7 | 250 |
| ▼ .local | | 0 | 156.7 | 250 |
| ▼ .call_fun | bivariate.R | 0 | 156.7 | 250 |
| ▼ <Anonymous> | bivariate.R | 0 | 156.7 | 250 |
| %*% | | 0 | 32.8 | 110 |
| ▶ .scale_n | SFEMethod-bivariate.R | 0 | 93.2 | 80 |
| is | SFEMethod-bivariate.R | 0 | 0.1 | 50 |
| t | | 0 | 30.7 | 10 |

g Moran bounds, adespatial

| Code | File | Memory (MB) | | Time (ms) |
|---|---|---|---|---|
| ▼ profvis::profvis | | -3703.4 | 6128.5 | 31180 |
| ▼ moran.bounds | | 0 | 5733.1 | 29700 |
| ▼ eigen | | 0 | 1918.2 | 23810 |
| ▼ isSymmetric.matrix | | 0 | 1660.8 | 1800 |
| all.equal.numeric | | 0 | 1149.1 | 900 |
| t.default | | 0 | 511.7 | 900 |
| ▼ bicenter.wt | | 0 | 3559.3 | 5070 |
| apply | | 0 | 1763.2 | 2110 |
| sweep | | 0 | 1024.8 | 1660 |
| t.default | | 0 | 771.3 | 1300 |
| t.default | | 0 | 255.4 | 630 |
| listw2mat | | 0 | 0.1 | 100 |
| range | | 0 | 0.2 | 10 |

h Moran bounds, Voyager

| | | | | |
|---|---|---|---|---|
| ▼ profvis::profvis | | 0 | 1276.9 | 2840 |
| ▼ moranBounds | | 0 | 1276.9 | 2090 |
| sweep | spatial-misc.R | 0 | 1021.3 | 1160 |
| t.default | | 0 | 255.4 | 660 |
| listw2mat | spatial-misc.R | 0 | 0.1 | 100 |
| W <- (W + t(W)) / 2 | spatial-misc.R | 0 | 0.0 | 100 |
| ▶ rowMeans | spatial-misc.R | 0 | 0.0 | 40 |
| colMeans | | 0 | 0.0 | 30 |

Supplementary Figure 2: Screenshots from profiling original implementations (left column) and the more efficient implementations in Voyager (right column). Both implementations were run on the same dataset unless otherwise noted. A) Most of the time is spent on re-finding distances between neighbors when using k nearest neighbors (knn) in spdep with inverse distance weighting (idw). B) Voyager avoids this time consuming step; the curve connects code in each implementation that finds the knn. C) Much of the time is spent on preprocessing the data frame input in the adespatial implementation of MULTISPATI PCA ("sweep" to scale and center, and "apply" to compute spatial lag). Also note that `eigen` was called twice. D) In the Voyager implementation, the direct input to the MULTISPATI function is a matrix. Much of the time was spent on scaling and centering the matrix (sweep) and computing the spatially weighted covariance matrix with matrix multiplication, which are much faster than the preprocessing steps in adespatial. For this dataset with around 1000 cells, time spent on partial eigen decomposition with RSpectra was negligible so it didn't show up in the profile. E) In spdep, much of the time was spent computing the spatially lagged values (lag.listw) and computing the sum of edge weights (lapply) when computing Lee's L for each pair of features. F) In Voyager, the listw spatial neighborhood graph is first converted to a sparse matrix so it's much faster to compute the sum of all non-zero entries and to compute the spatial lags for many features at once with matrix multiplication. For the same 347 genes, Voyager's implementation run on 5785 cells was much faster than spdep's implementation run on 105 cells in E. G) When finding Moran bounds with adespatial, most of the time was taken up by finding all the eigenvalues even though only the largest and smallest ones are needed. H) Most of the time and memory was spent on creating the double centered matrix (sweep, transpose, and etc.) while time spent finding the largest and smallest eigenvalues with RSpectra was negligible in Voyager's implementation of Moran bounds.

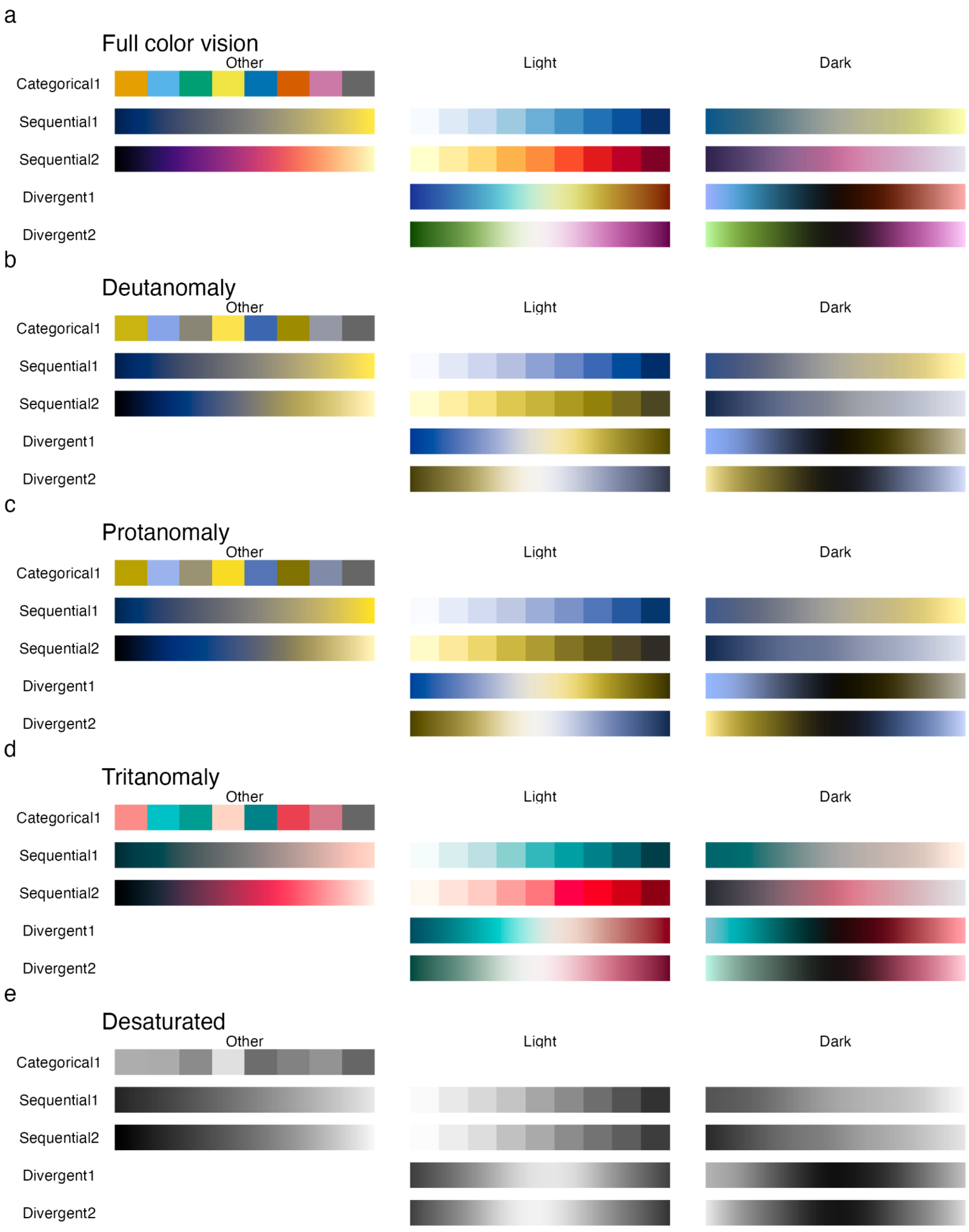

Supplementary Figure 3: Colorblind simulation of palettes in Voyager. A) All palettes used in the Voyager R package, full color vision. B) Deutanomaly perception of the palettes. C) Protanomaly perception of the palettes. D) Tritanomaly perception of the palettes. E)

Desatruated palettes. A dark theme is implemented to better visualize data with a fluorescent image in the background. The light and dark themes have different default palettes; in the light theme, darker color denotes higher values as if staining, while in the dark theme, lighter color denotes higher values as if glowing, so higher values stand out from the background. It is possible to simultaneously use two different palettes within the light or dark theme, such as to color Visium spots by one palette and cell segmentation from the same dataset with another in the same plot, but this should be used with caution. While the different palettes within one theme are chosen to avoid similar colors as much as possible, we do not suggest using two divergent palettes simultaneously, because doing so can distort color perception of either palette. We also do not suggest people with color vision deficiencies to use any two palettes simultaneously in one plot.

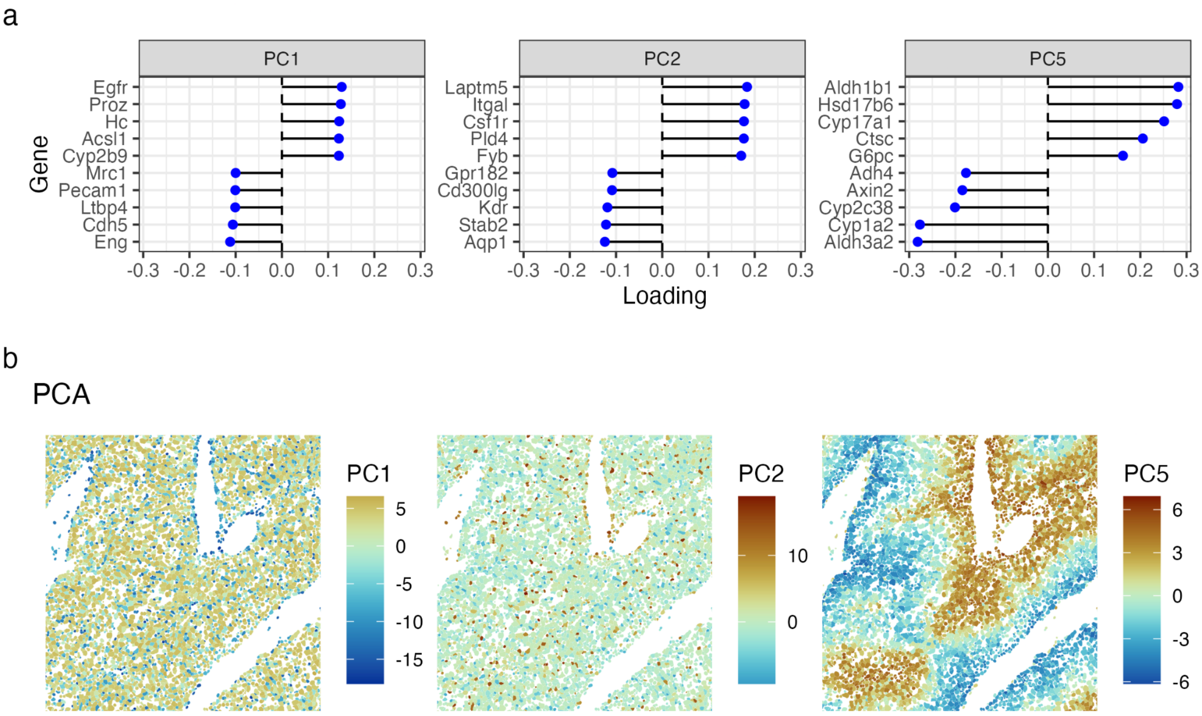

Supplementary Figure 4: Non-spatial PCA on mouse liver MERFISH dataset. A) Gene loadings of PCs 1, 2, and 5 from non-spatial PCA. B) MERFISH cell polygons colored by cell projections in PCs 1, 2, and 5.

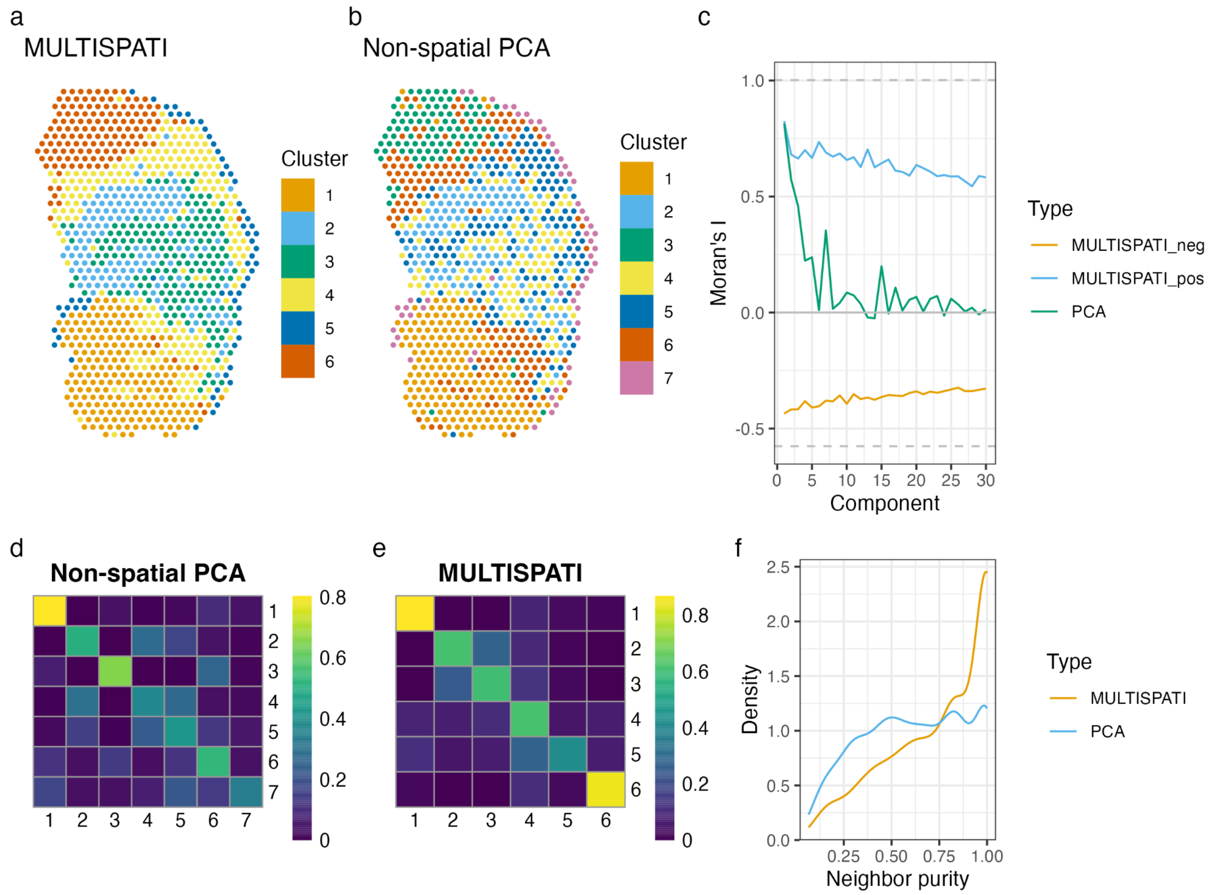

Supplementary Figure 5: Clustering with MULTISPATI and non-spatial PCA in mouse skeletal muscle Visium dataset. A) Leiden clusters with the first 30 positive MULTISPATI PCs in the mouse skeletal muscle Visium dataset. B) Leiden clusters with the first 30 non-spatial PC's. C) Moran's I of spot projection in each MULTISPATI and non-spatial PC. Moran's I decays sharply from non-spatial PC1 to PC5, but decays gradually for MULTISPATI PCs. D) Concordex heatmap for Leiden clusters based on non-spatial PCs and k nearest neighbor graph in histological space. This shows how well the clusters match the structure of the k nearest neighbor graph in histological space, although the graph in PCA space was used to generate the clusters instead. E) Concordex heatmap for clusters based on MULTISPATI PCs and k nearest neighbor graph in histological space, which shows a stronger diagonal and weaker off-diagonal entries indicating better spatial separation of clusters. The off-diagonal entries indicate proximity of spots from different clusters. F) Distribution of neighborhood purity of Leiden clusters based on non-spatial and MULTISPATI PCs and k nearest neighbor graph in histological space. MULTISPATI shows more spots with high neighborhood purity than non-spatial PCA.

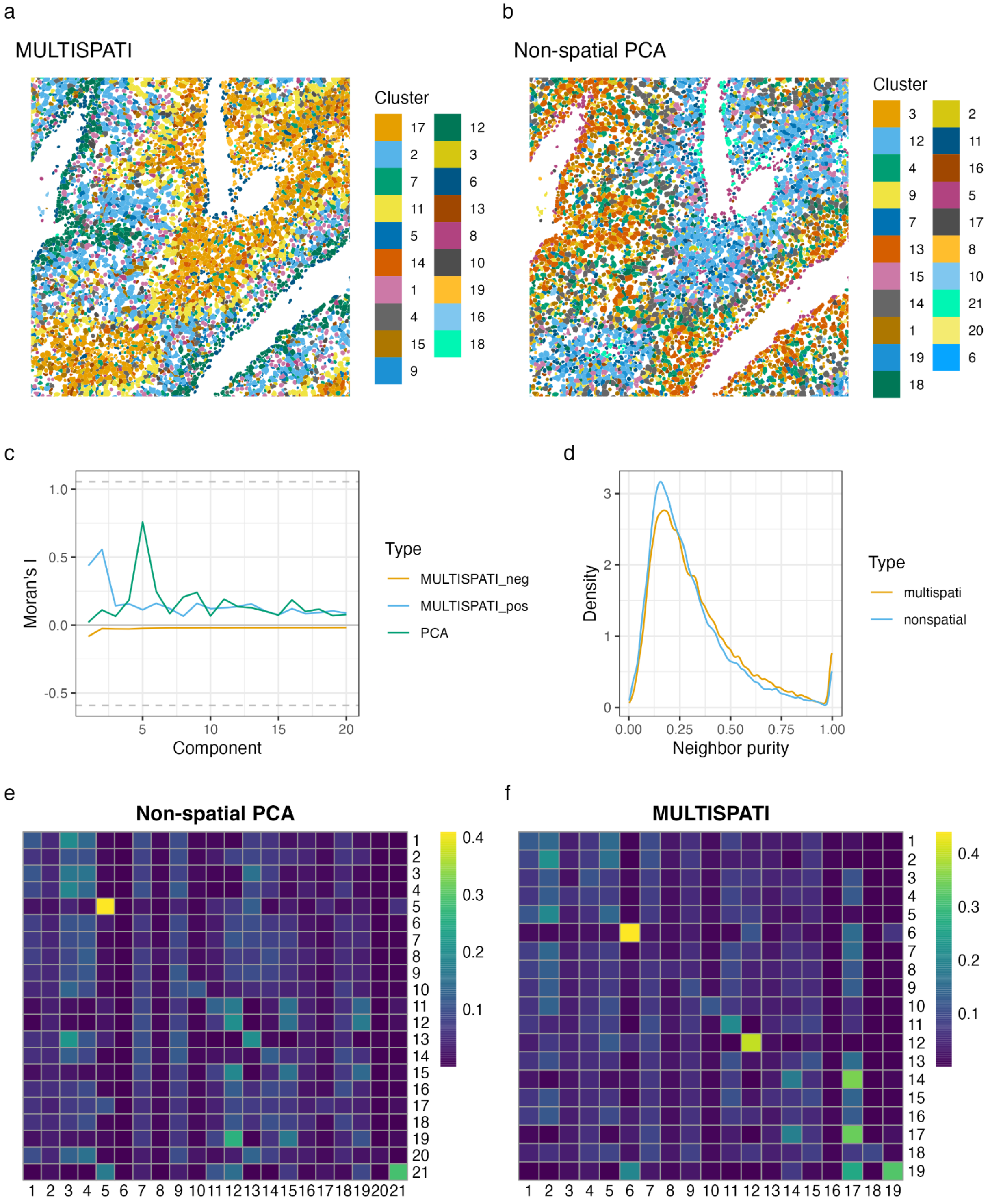

Supplementary Figure 6: Same as Supplementary Figure 5, but for mouse liver MERFISH dataset. The first 20 PCs are used for Leiden clustering. A-B) The clusters are visualized on a subset of the whole dataset while the clustering was performed on the whole dataset. Clusters are sorted by number of cells in the legend. In this case, from neighbor purity and Concordex, overall, MULTISPATI clusters are not markedly more spatially coherent than non-spatial PCA clusters. This may be because for this dataset, MULTISPATI does not produce more spatially coherent PCs than non-spatial PCA as shown in panel C. However, MULTISPATI gives a larger number of somewhat spatially coherent clusters than non-spatial PCA while the other clusters are not spatially coherent. Tools analyzing spatial colocalization of different cell types such as spicyR [12] can be used to find clusters based on colocalization of different cell types instead. Moran's I in this dataset should be interpreted differently from that in the Visium dataset, because in MERFISH, the spatial neighborhood graph used to compute Moran's I is based on single cells, while in Visium, the graph is based on spots which are aggregates of cells. Visium Moran's I indicates spatial autocorrelation of a longer length scale than MERFISH Moran's I.

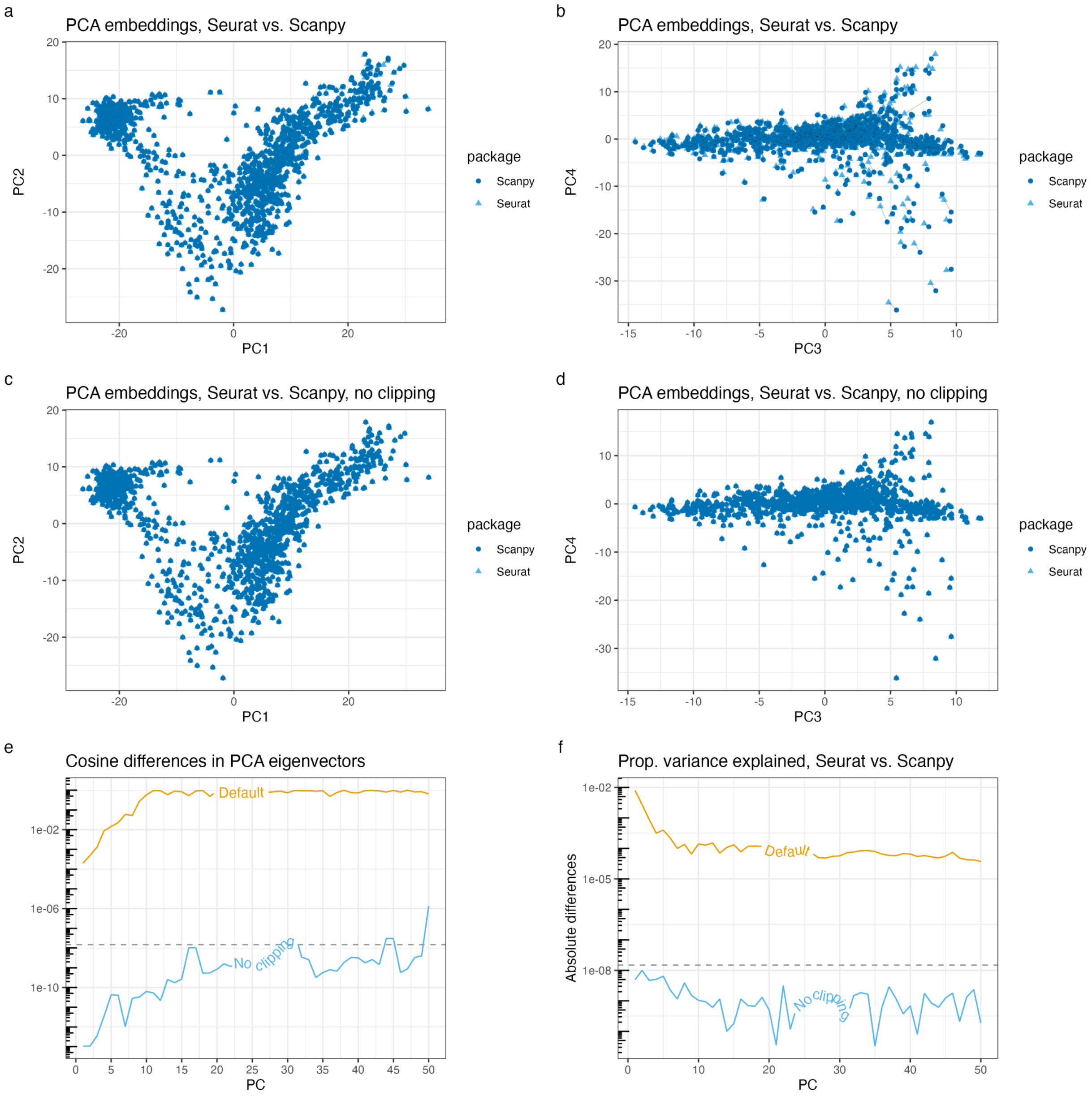

Supplementary Figure 7: Comparisons of PCA results from Seurat and scanpy using the same highly variable genes from Seurat. A) There's no visible difference between the Seurat and scanpy cell projections in the first 2 PCs. B) There are visible differences between Seurat and scanpy cell projections in PC3 and PC4. The lines connect corresponding cells from Seurat and scanpy. C-D) When not clipping the scaled data in Seurat, there's no visible difference between the Seurat and scanpy cell projections in the first 4 PCs. E) Cosine differences in each of the top 50 PCA eigenvectors between Seurat and scanpy. The dashed line is machine epsilon, or what can be accounted for by machine double precision. F) Absolute differences in proportion of variance explained by each PC in Seurat and scanpy. The differences are 5 orders of magnitude smaller without clipping than with default parameters. Without clipping, the differences are also within machine epsilon after the first two PCs, indicating that Seurat's clipping default which differs from that of scanpy is causing the different PCA results. E and F were made with the geomtextpath package v0.1.1.
